# A cellular hierarchy framework for understanding heterogeneity and predicting drug response in AML

**DOI:** 10.1101/2022.01.25.476266

**Authors:** Andy G.X. Zeng, Suraj Bansal, Liqing Jin, Amanda Mitchell, Weihsu Claire Chen, Hussein A. Abbas, Michelle Chan-Seng-Yue, Veronique Voisin, Peter van Galen, Anne Tierens, Meyling Cheok, Claude Preudhomme, Hervé Dombret, Naval Daver, P Andrew Futreal, Mark D. Minden, James A. Kennedy, Jean C.Y. Wang, John E. Dick

## Abstract

The treatment landscape of AML is evolving with promising therapies entering clinical translation, yet patient responses remain heterogeneous and biomarkers for tailoring treatment are lacking. To understand how disease heterogeneity links with therapy response, we determined the leukemia cell hierarchy make-up from bulk transcriptomes of over 1000 patients through deconvolution using single-cell reference profiles of leukemia stem, progenitor, and mature cell types. Leukemia hierarchy composition was associated with functional, genomic, and clinical properties and converged into four overall classes, spanning Primitive, Mature, GMP, and Intermediate. Critically, variation in hierarchy composition along the Primitive vs GMP or Primitive vs Mature axes were associated with response to chemotherapy or drug sensitivity profiles of targeted therapies, respectively. A 7-gene biomarker derived from the Primitive vs Mature axis was predictive of patient response to 105 investigational drugs. Thus, hierarchy composition constitutes a novel framework for understanding disease biology and advancing precision medicine in AML.

## Introduction

AML is a devastating disease characterized by extensive inter-patient and intra-patient heterogeneity. Poor outcomes are attributed to primary chemotherapy resistance and a high rate of relapse among patients who achieve remission, highlighting the inadequacy of standard chemotherapy as a means of curing AML for most patients. Recently, a wide range of promising new therapies targeting diverse cellular mechanisms have been approved or are progressing through clinical trials, offering alternatives to chemotherapy. However, patient responses to these new therapies are heterogeneous and we lack a reliable way to select the best therapy for each patient.

Historically, two distinct approaches have evolved for understanding heterogeneity in AML and informing therapy selection: a genomic model and a stem-cell model. The discovery of the Philadelphia chromosome in 1960 ^1^ sparked a series of cytogenetic studies that identified distinct cytogenetic drivers of AML. More recently, advances in genome sequencing have uncovered mutational drivers of AML and culminated in a prognostically informative genomic classification^2^. While this genomic model accounts for a major source of inter-patient heterogeneity, cells sharing the same driver mutation can exhibit functional differences ^3^. Moreover, while some driver mutations can be directly targeted by inhibitors, genomic profiling is limited in its ability to predict the benefit from therapies targeted to specific biological processes or signaling pathways.

The discovery of hematopoietic stem cells in 1961 and the development of quantitative assays to interrogate stem cell function ^4^ provided a basis for pioneering experiments that revealed that blasts within individual patients had functional differences in their cycling kinetics ^5–7^, differentiation state ^8–10^, and self-renewal capacity ^11–13^. Collectively, these studies conceptualized AML being sustained by rare leukemia stem cells (LSCs) ^14^, which were later formally identified through xenotransplantation studies ^15–17^. LSCs have since been shown to mediate relapse ^18^, and LSC-based gene expression stemness scores have emerged as predictors of outcomes following chemotherapy ^19–25^. While LSCs are an important therapeutic target, this model offers limited guidance around therapy selection. Overall, these genomic and stem-cell models provide complementary insight into AML heterogeneity, yet neither model alone is sufficient to guide therapy selection, particularly for newer agents. A new approach for personalized therapy selection that integrates the genomic and stem cell models is needed.

Cancer has long been recognized as a caricature of normal tissue development ^26, 27^. AML is one of the best-studied cancer systems wherein leukemic cells are organized into a hierarchy resembling normal blood development. Cellular hierarchies in AML can be distorted in different ways, depending on genetic alterations and cell of origin. For example, a strong differentiation block arising in a stem cell may result in a shallow, stem cell-dominant hierarchy. In other cases considerable - albeit aberrant - differentiation may occur resulting in a steep hierarchy wherein rare LSCs generate a bulk blast population with mature myeloid features. In this way, the cellular composition of each patient’s leukemic hierarchy likely reflects the functional impact of specific mutations on the disease-sustaining LSCs. Thus, interrogation of leukemic hierarchies may provide an opportunity to potentially integrate features of the genetic and stem cell models of AML ^28^. Single-cell RNA-sequencing (scRNA-seq) has emerged as a powerful tool for dissecting cellular hierarchies ^29, 30^, however prohibitive costs restrict these studies to a limited number of patients. Without measuring these cellular hierarchies at scale in large clinical datasets, the relationship of hierarchy composition to therapy response remains unknown.

Here, we were able to characterize the cellular hierarchies of >1000 AML patients through gene expression deconvolution on bulk AML transcriptomes using single-cell reference profiles of distinct AML stem, progenitor, and mature types. This approach to characterizing AML heterogeneity enabled integration of both the genomic and functional models of AML resulting in a novel framework for understanding disease biology and predicting drug response.

## Results

### Single-cell characterization of leukemia stem and progenitor populations

As a first step to uncover the organization of cellular hierarchies in AML, we re-analyzed the scRNA-seq data of 13,653 cells from 12 AML patients at diagnosis ^29^ with a focus on primitive stem and progenitor blast populations (henceforth Leukemia Stem and Progenitor Cells, LSPCs). Using Self-Assembling Manifolds (SAM), an unsupervised approach to prioritize biologically relevant features among relatively homogenous cells ^31^, we previously identified two transcriptomic populations of normal human HSC: a deeply quiescent population with low transcriptome diversity (Non-Primed) and another residing in a shallower state of quiescence with higher CDK6 expression (Cycle-Primed) ^32^. We applied SAM to analyze LSPCs and identified three distinct populations shared across the 12 patients (Fig. 1A, Fig. S1A-C). One population had low transcriptome diversity and was enriched for core LSC programs but appeared otherwise inactive (Fig. S1D). We named this population Quiescent LSPC. The second population was enriched for CDK6 expression and targets of the cell cycle regulator E2F3 suggestive of cell cycle priming (Fig. S1E), as well as inflammatory signatures suggestive of priming for myeloid differentiation ^33^ (Fig. S1H). We named this population Primed LSPC. The third population exhibited enrichment for CTCF targets suggestive of stem cell activation ^34^ and broad enrichment of E2F targets indicating cell cycle progression, with 40% of cells classified as cycling (Fig. 1B, S1F-G, I). We named this third population Cycling LSPC. The existence of distinct cellular states provides a molecular basis for the known functional heterogeneity that is found within the LSC compartment ^35^. These new classes of Quiescent, Primed, and Cycling LSPC led to higher classification performance compared to the prior ‘HSC-like’ and ‘Progenitor-like’ classification from van Galen *et al* ^29^ (weighted accuracy: 0.93 vs 0.73, respectively; Fig. S1J). We combined these new LSPC classes with the existing classification of more committed blasts by van Galen *et al* (GMP-like blasts resembling Granulocyte-Monocyte Progenitors, ProMono-like blasts resembling promonocytes, Mono-like blasts resembling monocytes, and cDC-like blasts resembling conventional Dendritic cells) to constitute a map of common leukemic blast states shared across these 12 AML patients, with each leukemic state having distinct molecular properties.

**Figure 1.**
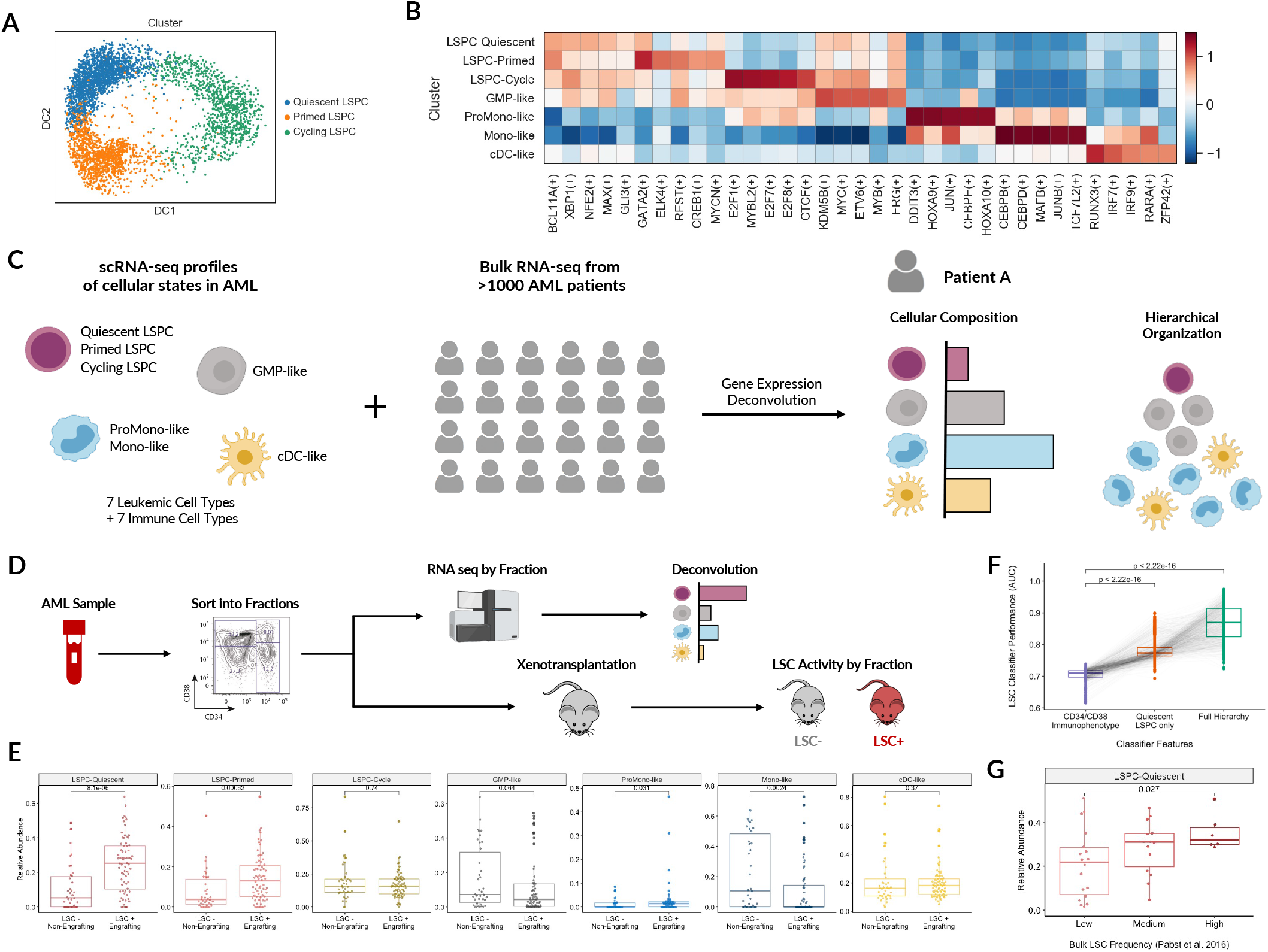
Functional significance of LSPC populations from single-cell RNA-seq. A) Diffusion map of 4163 AML Leukemia Stem and Progenitor Cells (LSPCs) using feature weights from Self-Assembling Manifolds (SAM). B) Transcription Factor regulon activity in each leukemic cell type, inferred through PySCENIC. Transcription factor regulon enrichment scores were scaled and the top five regulons for each cell type are depicted. C) Schematic of deconvolution approach using reference signatures from single-cell RNA-seq populations. D) Experimental design for evaluating the relationship between AML cell states from scRNA-seq and functional LSC potential. 111 sorted AML fractions previously evaluated for functional LSC activity through xenotransplantation in Ng et al (2016) were subject to RNA sequencing and gene expression deconvolution. The relative abundance of each leukemic population was subsequently compared across LSC+ (engrafting) and LSC- (non-engrafting) fractions. E) Enrichment of leukemic cell types across LSC+ (engrafting) and LSC- (non-engrafting) AML fractions. Relative abundances of each cell type were compared through a Wilcoxon rank-sum test. F) Model performance (AUC) of Random Forest classifiers predicting functional LSC activity in sorted AML fractions. Classifiers were trained and evaluated through 5-fold nested cross-validation with 1000 repeats, as outlined in Figure S3A. Three types of classifiers were trained, each using different features to predict LSC activity: (1) using the CD34/CD38 immunophenotype of each fraction, (2) using the relative abundance of Quiescent LSPC alone, and (3) using the relative abundance of all leukemic populations spanning the full AML hierarchy. AUC values are paired by iteration, wherein sample order and cross-validation splits were identical for each classifier. Comparisons were performed using a Wilcoxon signed-rank test. G) Relative abundance of Quiescent LSPC in patient samples with low, medium, and high bulk LSC frequencies, as defined by Pabst et al (2016). Comparisons were performed using a Wilcoxon rank-sum test.

### Deconvolution of constituent cell populations in AML

We next sought to understand how these defined AML cell populations and the hierarchies into which they are organized relate to functional, biological, and clinical properties of AML. To study this at scale, we employed gene expression deconvolution to infer the leukemic hierarchy composition from bulk AML transcriptomes (Fig. 1C). We performed benchmarking analysis of multiple scRNA-seq-based deconvolution methods and identified CIBERSORTx as the highest-performing approach in the context of AML (Supplementary Note 1, Fig. S2). Additionally, we confirmed that deconvolution with the new LSPC classification of primitive AML cells improves discrimination of clinical and biological phenotypes compared to the prior HSC-like and Prog-like classification (Supplementary Note 2, Fig. S3). Thus, subsequent deconvolution was performed through CIBERSORTx using single-cell transcriptomes from seven leukemic cell types (Quiescent LSPC, Primed LSPC, Cycling LSPC, GMP-like, ProMono-like, Mono-like, cDC-like) and seven non-leukemic immune cell types (Natural Killer, Naive T, CD8+ T, B, Plasma, Monocytes, and cDCs) as a reference.

### Quiescent LSPC abundance is associated with functional LSC activity

We first sought to determine whether any of our newly-defined LSPC cellular states were associated with LSC activity. The LSC state is functionally defined by whether a leukemic cell can initiate leukemia *in vivo* ^36^. We thus performed RNA-seq on 111 AML fractions previously evaluated by microarray and where LSC activity was determined through xenotransplantation ^23^ and applied deconvolution to determine the cell type composition of each fraction (Fig. 1D). LSC-positive fractions were highly enriched for Quiescent LSPC (p = 8e-6) and Primed LSPC (p = 6e-4) but not Cycling LSPC (p = 0.74) (Fig. 1E). Conversely, LSC-negative fractions were highly enriched for Mono-like blasts (p = 2e-3) (Fig. 1E). Given that immunophenotype does not consistently predict LSC activity ^20, 23, 37^, we compared deconvolution against immunophenotype by training classifiers to predict LSC activity in AML fractions based on cell type composition versus CD34/CD38 status. Classifiers trained on immunophenotype were consistently outperformed by those trained on leukemia cell composition (median AUCs = 0.71 vs 0.86, p < 2e-16) and were even outperformed by models trained from Quiescent LSPC abundance as a single variable (median AUCs = 0.71 vs 0.77, p < 2e-16) (Fig. 1F, Table S3). Finally, we found Quiescent LSPC to be associated with high LSC frequency in an independent dataset of bulk AML samples assessed through limiting dilution analysis ^38^ (Fig. 1G). Collectively, these findings establish a new link between transcriptomic LSPC states and functionally defined LSC at the apex of the hierarchy, suggesting that LSC activity can be inferred through deconvolution of patient hierarchies.

### AML hierarchy composition is associated with clinical and genomic properties

The differentiation properties of the LSCs sustaining each patient’s AML are reflected in the cellular composition of the hierarchies that they generate. To examine how these hierarchies vary across patient samples and how they relate to molecular and clinical features of AML, we applied our deconvolution approach to infer the abundance of 7 leukemic cell types as well as 7 non-leukemic immune populations (described above) within 864 patient samples from the TCGA ^39^, BEAT-AML ^40^, and Leucegene cohorts ^41^. Clustering patients based on the composition of their leukemia cell hierarchies revealed four distinct subtypes: Primitive (shallow hierarchy, LSPC-enriched), Mature (steep hierarchy, enriched for mature Mono-like and cDC-like blasts), GMP (dominated by GMP-like blasts), and Intermediate (balanced distribution) (Fig. 2A-C). Hierarchy composition was associated with multiple biological and clinical parameters including age at diagnosis, WBC differential counts, and FAB class (Fig. S4A-B). We focused on cytogenetic and mutational correlates in order to understand the cellular states and hierarchies generated by common genetic drivers of AML.

**Figure 2.**
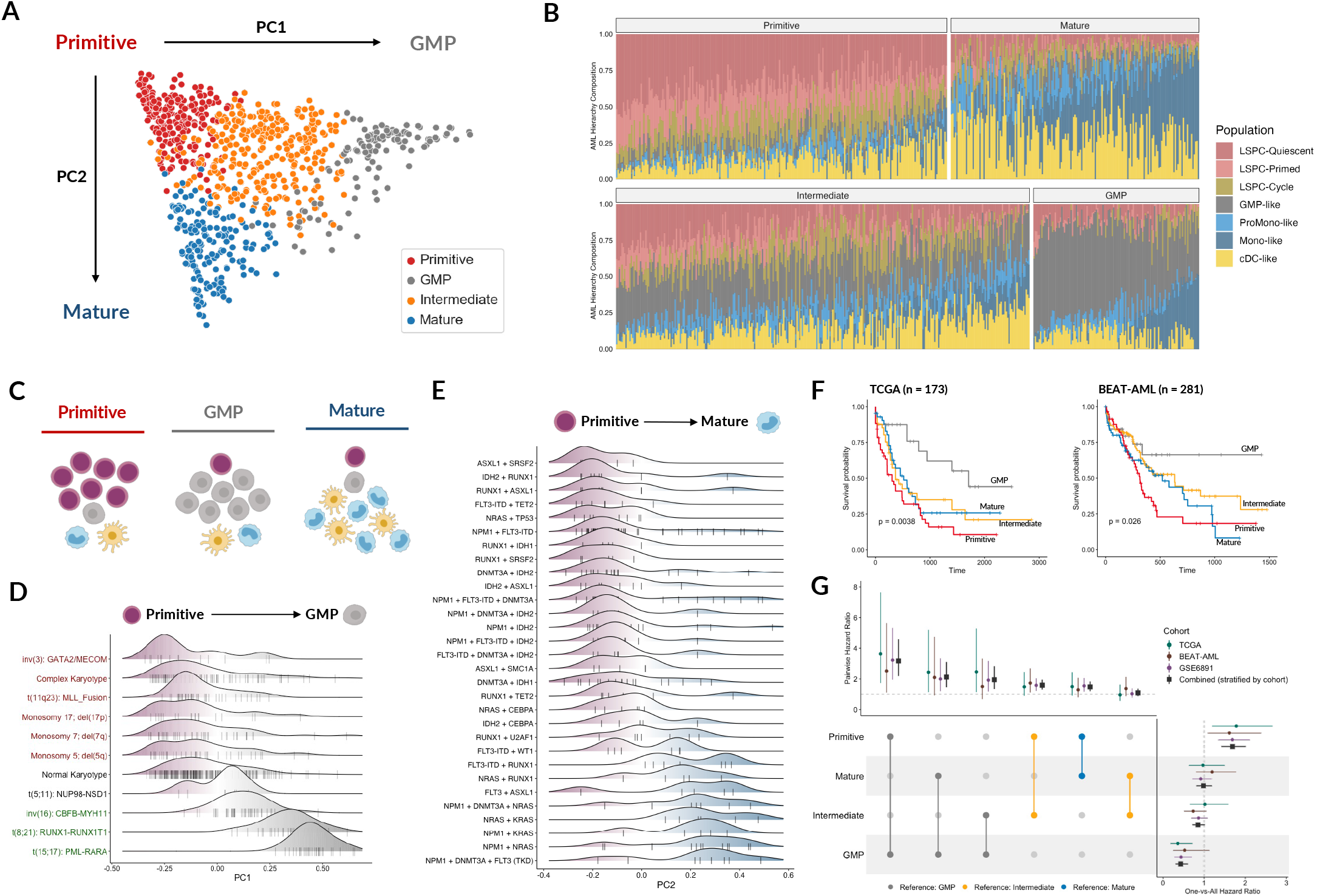
AML hierarchy composition correlates with genomics and survival. A) Principal component analysis of 864 AML patients from the TCGA, BEAT-AML, and Leucegene cohorts based on the composition of their cellular hierarchy. B) Relative abundance of each leukemic cell type in each patient. Each bar represents an individual patient and the distribution of colors throughout each bar represents the distribution of leukemic cell populations within their leukemic hierarchy. C) Depictions of the cellular organization of Primitive, GMP, and Mature hierarchies. D) Density plots depicting cytogenetic groups along the Primitive vs GMP axis (PC1). Cytogenetic alterations are coloured by prognostic significance, wherein red indicates adverse prognosis while green indicates favorable prognosis. E) Density plots depicting common driver mutation combinations along the Primitive vs Mature axis (PC2). F) Overall survival outcomes of AML hierarchy subtypes in the TCGA and BEAT-AML cohorts. Differences in survival across all subtypes were evaluated through a log-rank test. G) Univariate and pairwise hazard ratios for each AML hierarchy subtype across three patient cohorts. Univariate hazard ratios for each subtype and pairwise hazard ratios between subtypes are depicted alongside their 95% confidence intervals. Pairwise comparisons are colored based on the reference subtype, which is always positioned lower than the query cluster. Combined hazard ratios, obtained by pooling individual patient outcomes and performing Cox proportional hazards regression stratified by cohort, are also depicted by black squares alongside the hazard ratios derived from each individual cohort.

Patient hierarchies were separated along two principal components: PC1, spanning a continuum from Primitive to GMP (35% of variance), and PC2, spanning Primitive to Mature (28% of variance) (Fig. 2A). Hierarchies generated by cytogenetic alterations primarily separated along the Primitive vs GMP axis (PC1) (Fig. 2D; Fig. S4E), with adverse cytogenetic alterations generating Primitive hierarchies and favorable cytogenetic alterations generating GMP-dominant hierarchies (Fig. 2D). Cellular hierarchies generated by genetic mutations and their combinations primarily separated along the Primitive vs Mature axis (PC2), reflecting their impact on the extent of AML differentiation (Fig. 2E; Fig S4C-D). Notably, different mutations in the same gene could have different consequences on the resulting hierarchies. For example, DNMT3A R882 mutations were associated with more mature disease than other DNMT3A mutations (Fig. S4F-G) suggesting that DNMT3A R882 may be more permissive of AML differentiation compared to other DNMT3A mutations. Collectively, these data demonstrate that describing AML inter-patient heterogeneity based on hierarchy composition can capture and integrate both genomic and stem cell models of AML heterogeneity.

### Primitive vs GMP axis captures patient prognosis

In line with the observed associations with favorable and adverse cytogenetics, patients with different hierarchy subtypes also differed in their survival outcomes, with Primitive hierarchies associated with the worst outcomes and GMP-dominant hierarchies associated with the best outcomes (Fig. 2F-G, Table S6). We validated these findings using microarray data from a cohort of genetically diverse adult AML patients (GSE6891 ^42, 43^; Fig. S5A-B) as well as a cohort of pediatric AML patients (TARGET-AML ^44^; Fig. S5C-D). To identify the leukemic cell types linked to patient survival, we performed regularized cox regression on the TCGA and BEAT-AML datasets using leukemia cell type abundances. Quiescent LSPC and Cycling LSPC abundance were predictive of adverse outcomes (coefficients: 0.34 and 0.72, respectively) and GMP-like abundance was predictive of favorable outcomes (coefficient: -1.54). Strikingly, the composite survival score that included all three of these populations was highly anti-correlated with PC1 (r = -0.99, Fig. S5E). Indeed, PC1 was highly associated with survival outcomes in TCGA, BEAT-AML, and GSE6891 cohorts (Fig. S5F), and retained significance in a multivariate meta-analysis incorporating all three datasets (p = 0.007, Table S7). Moreover, both pediatric and adult AML patients who did not achieve complete remission following induction chemotherapy had higher Quiescent LSPC abundance and lower GMP-like abundance compared to patients who achieved remission (Fig. S5G). In contrast, the Primitive vs Mature axis was not associated with patient survival (p = 0.412, Table S7).

We reasoned that the biology underlying the Primitive vs GMP axis may also underlie part of the variation captured by existing prognostic scores in AML. Indeed, when we considered four recent prognostic gene expression scores for AML (LSC17 ^23^, APS ^45^, 3-Gene ^46^, CODEG22 ^47^) we found convergence across all four scores wherein patients with high scores had high Quiescent LSPC abundance and low GMP-like abundance (Fig. S5H-I). Finally, unbiased analysis of the association between individual genes with survival outcomes revealed that genes associated with shorter survival were enriched for HSC-specific programs while genes associated with longer survival were enriched for GMP-specific programs (Fig. S5J).

Although prior studies implicated primitiveness as reflecting poor outcomes, our data reveal more complexity. Rather, our data suggest that the biological distinction between stem cells and GMP progenitors underlie prognosis in AML rather than stemness properties alone. Thus our data argue that existing prognostic AML scores are linked to this specific axis pointing to the clinical importance of the biological properties that determine hierarchy composition.

### Hierarchy composition changes between diagnosis and relapse

Given the associations observed between hierarchy composition and clinical outcomes in AML, we asked whether the composition of these hierarchies evolve over the course of disease. To understand how the AML hierarchies change throughout disease progression, we deconvoluted 44 pairs of AML samples collected at diagnosis and relapse following induction chemotherapy from four independent cohorts ^18, 48–50^ (Fig. 3A, S6A). At diagnosis, patients presented with diverse hierarchy compositions, yet by relapse most were classified as Primitive (Fig. 3B) with significant expansion of total LSPC populations (p = 1e-8) and, in particular, Quiescent LSPC (p = 9e-6, Fig. 3C). To validate this finding at the single-cell level, we analyzed scRNA-seq data from 8 relapsed AML patients ^51^ and observed uniformly higher LSPC abundance as compared to scRNA-seq data from 12 diagnostic AML samples ^29^ (Fig. 3D, S6B).

**Figure 3.**
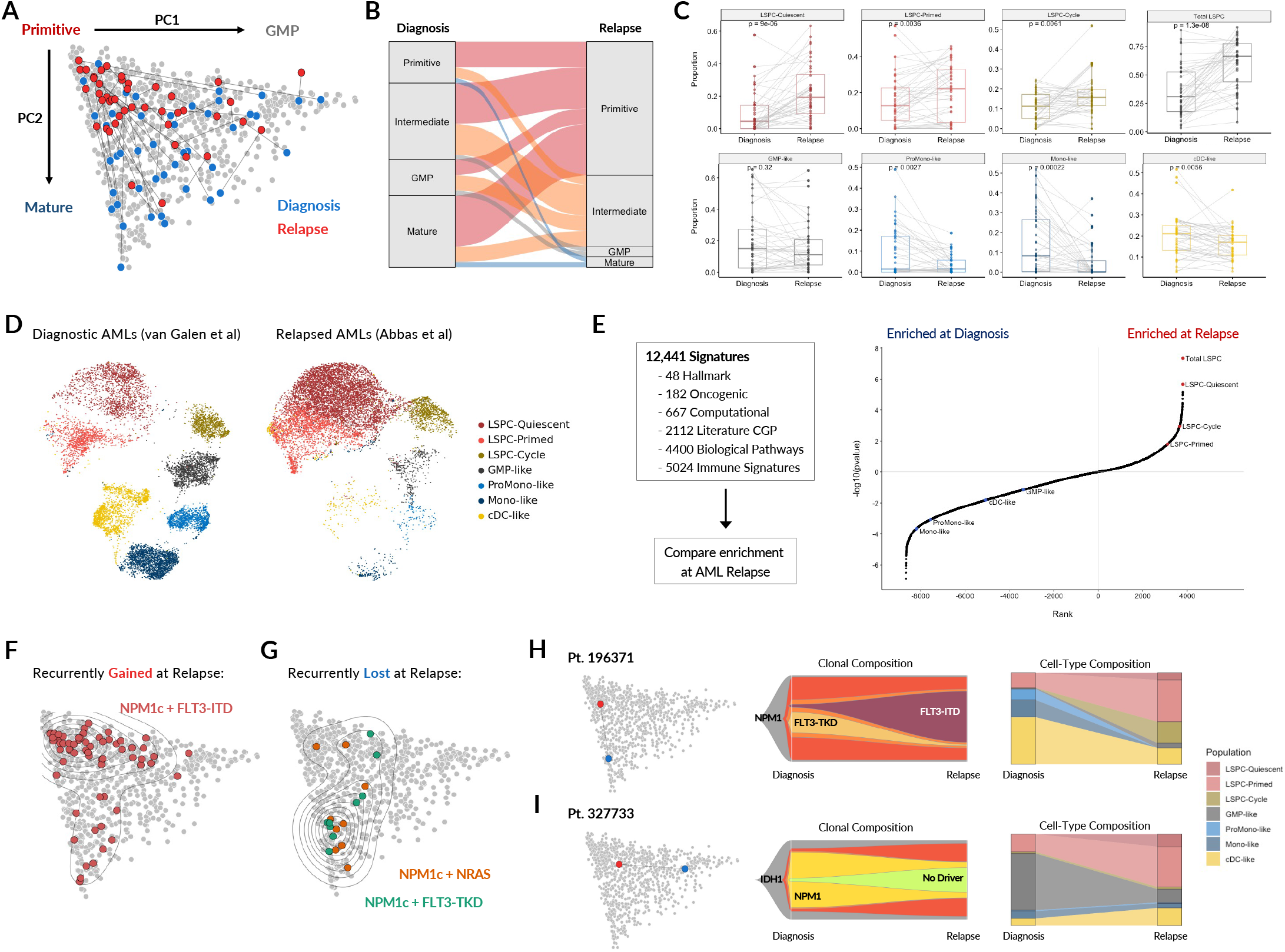
Transitions in hierarchy composition from diagnosis to relapse. A) Transitions in hierarchy composition from diagnosis to relapse in 44 paired AML samples from four independent cohorts. B) Alluvial diagram depicting hierarchy subtype distribution from diagnosis to relapse. The width of each band reflects the number of patients transitioning from one subtype to another from diagnosis to relapse. C) Changes in leukemic cell type abundance from diagnosis to relapse, including changes in total LSPC abundance. Significance was evaluated using a Wilcoxon signed-rank test. D) Single-cell RNA-seq of diagnostic AML from van Galen et al (2019) compared to relapsed AML samples from Abbas et al (2021), classified, downsampled to 10,000 cells, and projected onto a common embedding using scArches with scANVI. E) Benchmarking the significance and magnitude of changes in hierarchy composition from diagnosis to relapse against 12,441 signatures from MSigDB. The rank and significance of enrichment for each leukemic cell population as well as 12,441 signatures from diagnosis to relapse are shown, with the Y-axis depicting the absolute value of the log10(pvalue) in the direction of the enrichment (positive for relapse, negative for diagnosis). P-values were derived from paired t-tests; non-parametric Wilcoxon signed-rank tests were also performed to ensure that results were comparable. F) Patients with NPM1 + FLT3-ITD (recurrently acquired at relapse) G) Patients with NPM1c + NRAS, and NPM1c + FLT3-TKD (recurrently lost at relapse). NRAS and FLT3-TKD mutations were filtered at a 0.25 VAF cutoff. H-I) Changes in clonal and cell type composition from diagnosis to relapse. These are depicted for a patient with concordant shifts in both clonal composition and cell type composition (H) as well as for a patient with dramatic changes in cell type composition with minimal detected changes in clonal composition with respect to known driver mutations (I).

While LSPC expansion at relapse is in line with prior functional xenotransplantation studies, the consistency and magnitude of this phenotype were unexpected, with LSPC expansion observed in 89% of patients at relapse (39 of 44 pairs). Furthermore, all 5 patients for whom LSPC abundance decreased at relapse already had high LSPC abundance at diagnosis (median 78.9%, compared to 27.3% for other patients). To benchmark this finding, we also evaluated 12,441 biological signatures from MSigDB spanning biological pathways, immune processes, and cancer/AML-specific gene sets, and found enrichment in total LSPC at relapse to be two orders of magnitude more significant than the top-ranked signature from MSigDB (Fig 3E). Indeed, classifiers trained on the abundance of LSPC populations were able to achieve near-perfect performance in classifying paired diagnosis and relapse samples (median AUC = 0.96; Table S8).

These changes in cellular composition from diagnosis to relapse can help contextualize patterns of clonal evolution in AML. For example, FLT3-ITD alterations are recurrently gained at relapse while NRAS and FLT3-TKD alterations are recurrently lost at relapse in NPM1-mutant AMLs ^52^. Indeed, FLT3-ITD with NPM1c generated Primitive hierarchies (Fig. 3F), while NRAS or FLT3-TKD with NPM1c generated Mature hierarchies (Fig. 3G). For a subset of the patients we analyzed, changes in composition from diagnosis to relapse were concordant with patterns of clonal evolution (Fig. 3H, Fig. S6C-F). In other cases, shifts in composition occurred in the absence of clear genetic changes, potentially due to non-genetic modes of evolution (Fig. 3I). Together, our findings establish LSPC population expansion as a common hallmark across diverse evolutionary paths to AML relapse following chemotherapy.

### Primitive vs Mature axis predicts sensitivity to investigational drugs

Having shown that survival outcomes following chemotherapy are tied to hierarchy composition (i.e. Primitive vs GMP axis), we asked whether AML samples with different cellular compositions were vulnerable to newer investigational therapies. Ex vivo drug sensitivity data from two public datasets ^40, 53^ were integrated with cell type abundance to generate drug sensitivity profiles for each leukemic cell type (Fig. 4A). This revealed large differences in drug responses between primitive blasts and mature blasts, with separation of drug responses occurring primarily along the Primitive vs Mature axis, in which PC2 significantly correlated (FDR < 0.05) with response to 37 drugs in the BEAT-AML screen and 64 drugs in a separate screen from Lee et al (Total = 101; Fig. 4B, S7A). By contrast, the Primitive vs GMP axis (PC1) was not associated with sensitivity to any drug from either screen (Fig. S7A).

**Figure 4.**
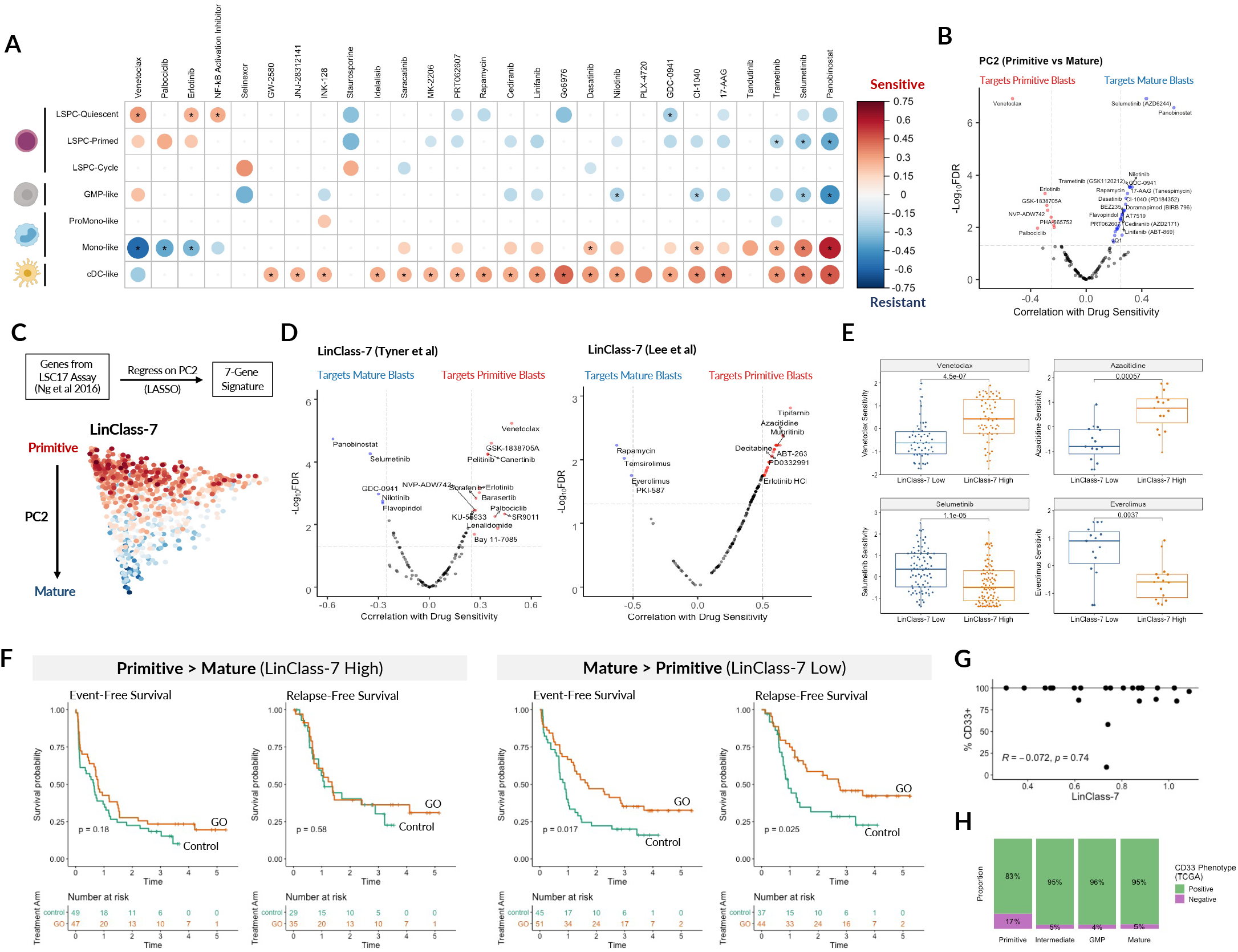
AML Hierarchy composition as a determinant of targeted therapy response. A) Correlation between cell type abundance and *ex vivo* drug sensitivity (-AUC) across 202 diagnostic patient samples in BEAT-AML, wherein color and size represent the direction and magnitude of the correlation. Only correlations with p < 0.05 are depicted, those with q < 0.05 are marked with an asterisk. B) Volcano plot of correlations between the Primitive vs Mature axis (PC2) and *ex vivo* drug sensitivities from the BEAT-AML screen, identifying drugs that preferentially target either primitive or mature AML blasts. C) LinClass-7 (trained on PC2) captures the Primitive vs Mature axis. D) Correlation of LinClass-7 identifies drugs targeting either primitive blasts or mature blasts from BEAT-AML (Tyner et al, 2018; n = 202) as well as a separate primary AML drug screen (Lee et al, 2019; n = 30). E) Venetoclax and Azacytidine target primitive AML blasts (LinClass-7 high), MEK and mTOR inhibition targets mature AML blasts (LinClass-7 low). F) Subgroup analysis of the ALFA-0701 trial, evaluating Gemtuzumab-Ozogamicin (GO), a drug-conjugated antibody targeting CD33 in AML. Event-free survival and relapse-free survival of control patients (Daunorubicin + Cytarabine) compared to GO patients (Daunorubicin + Cytarabine + GO), stratified by LinClass-7 score into LinClass-7 High (Primitive > Mature) and LinClass-7 Low (Mature > Primitive). G) Lack of correlation between LinClass-7 and surface CD33 levels, evaluated across 23 Toronto PMH AML patients for which both RNA-seq and clinical flow information was available. H) CD33 surface marker phenotype of TCGA patients, demonstrating very high rates of CD33 positivity among AML patients regardless of hierarchy composition.

The large number of drugs for which sensitivity was associated with the Primitive vs Mature axis suggested that this axis constituted the primary source of variation underlying ex vivo sensitivity to investigational drugs in AML. To test this hypothesis, we performed unsupervised clustering of patient samples from Lee et al ^53^ based on their ex vivo sensitivity to 159 drugs and identified two patient clusters with global differences in their drug sensitivity profiles (Fig. S7B). Differential expression analysis revealed that one cluster was highly enriched for primitive HSC programs while the other was enriched for mature myeloid programs (Fig. S7C), demonstrating that the Primitive vs Mature axis captures fundamental differences in drug sensitivity profiles among AML patients.

### Hierarchy-based gene expression scores predict drug sensitivity in AML

As a first step to translate the association between the Primitive vs Mature axis and an individual’s response to a specific drug into the clinic, we sought to capture this axis through simple gene expression scores. As a proof of concept, we turned to the LSC17 score, for which a CAP/CLIA-certified clinical assay has been developed on the NanoString platform ^23, 54^. Given that the LSC17 score was associated with leukemic hierarchy composition (Fig. S5H-I), we reasoned that deriving a sub-score from these 17 genes to estimate PC2 may provide a rapidly deployable tool to inform therapy selection using data from the existing LSC17 assay. We thus retrained the LSC17 genes on PC2 through LASSO regression with leave-one-out cross-validation to identify a 7-gene lineage classification sub-score (hereafter LinClass-7) (Fig. 4C). LinClass-7 correlated well with PC2 (|r| = 0.82) in the validation cohort and was significantly associated with sensitivity to 33 drugs from BEAT-AML as well as 72 drugs from Lee et al (Total = 105; Fig. 4D), each of which targeted either primitive blasts (e.g. Venetoclax, Azacytidine, Mubritinib) or mature blasts (e.g. MEK/mTOR inhibitors) (Fig. 4E). Importantly, neither LSC17 nor the other prognostic AML scores ^45–47^ were significantly associated with drug sensitivity (Fig. S7D) and none effectively stratified patients by sensitivity to clinically relevant agents such as Venetoclax and Azacitidine (Fig. S7E).

To examine the clinical relevance of these Primitive vs Mature scores, we turned to gene expression data from the pivotal ALFA-0701 adult AML clinical trial of low fractionated doses of Gemtuzumab Ozogamicin (GO) in combination with standard chemotherapy ^55, 56^. We asked whether the LinClass-7 score could predict clinical benefit from GO treatment. In the ALFA-0701 trial, the addition of GO to standard chemotherapy conferred significant benefits in event-free survival and relapse-free survival (EFS HR = 0.64 [0.47 - 0.89], p = 0.008; RFS HR = 0.65 [0.43 - 0.99], p = 0.045; Table S9), but differences in overall survival were not significant at the final time of follow-up. LinClass7 effectively separated responders from non-responders: GO treatment led to significantly longer event-free survival and relapse-free survival (EFS HR = 0.57 [0.35 - 0.91], p = 0.018; RFS HR = 0.53 [0.30 - 0.93], p = 0.028; Table S9) for patients with LinClass-7 low (Primitive > Mature) AML, although this association did not extend to overall survival. In contrast, patients with LinClass-7 high (Mature > Primitive) AML derived no significant survival benefit from GO (Fig. 4F, Table S9). Importantly, we observed no association between surface levels of CD33 (the molecular target of GO) with either LinClass-7 (Fig. 4G) or PC2 (Fig. S8B). In fact, CD33 levels appeared to be unassociated with hierarchy composition: most patients were CD33 positive regardless of their hierarchy subtype (Fig. 4H). The LSC17 score has also been shown to predict clinical benefit from GO treatment ^23^. Our analysis shows that the LSC17 and LinClass-7 scores captured different subsets of patients and further subgroup analysis revealed that only patients that had low scores for both LinClass-7 and LSC17 derived clinical benefit from GO treatment (EFS HR = 0.44 [0.23 - 0.85], p = 0.014; RFS HR = 0.35 [0.16 - 0.78], p = 0.009; Fig. S8C; Table S9), demonstrating complementarity between LSC17 and LinClass-7 in the prediction of clinical benefit from GO.

In the context of adult AML, these analyses point to the utility of LinClass-7 as a companion score to LSC17, enabling prediction of response to an array of current and investigational drugs. Importantly, the distribution of LSC17 and LinClass-7 scores across patient samples also loosely recapitulate the primary axes of variation in hierarchy composition, separating Primitive, GMP, and Mature AMLs (Fig. S9A). Thus the LSC17 and LinClass-7 scores, measurable through the same CAP/CLIA-certified clinical assay ^54^ (Fig S9B), allow for both prognostic and predictive stratification of patient samples while also providing salient information on patient hierarchy composition.

Although adult and pediatric AML are molecularly distinct diseases ^44, 57^, the Primitive vs Mature axis captured through PC2 was also able to predict clinical benefit from GO among pediatric AML patients in the TARGET-AML retrospective cohort ^44^. Patients with high PC2 (Mature > Primitive) experienced longer overall and event-free survival outcomes with GO treatment compared to those that did not receive GO treatment (OS HR = 0.51 [0.30 - 0.89], p = 0.017; EFS HR = 0.58 [0.37 - 0.90], p = 0.017; Table S11). In contrast, GO treatment did not influence survival outcomes of patients with low PC2 (Primitive > Mature) (OS HR = 1.20 [0.68 - 2.00], p = 0.553; EFS HR = 0.97 [0.62 - 1.50], p = 0.878) (Fig. S8E-F, Table S11). However, in the context of pediatric AML we observed lower correlation between LinClass-7 and PC2 (r = 0.51) and found that prediction of clinical benefit from GO treatment in pediatric AML did not extend to either LinClass-7 or LSC17 (Fig. S87E-F, Table S11).

Given this, we asked whether a Primitive vs Mature score not constrained by the LSC17 genes could accurately recapitulate PC2 in both adult and pediatric AML. Starting with the top 100 PC2-associated genes, we trained a 34-gene score (termed PC2-34) which was highly correlated with PC2 (|r| = 0.95 in the validation cohort). PC2-34 performed similarly to LinClass-7 in capturing drug sensitivity in the BEAT-AML (48 drugs at FDR < 0.05) and Lee et al (82 drugs at FDR < 0.05) screens (Fig. S7D-E). PC2-34 also predicted clinical benefit from GO treatment (Fig. S8A, Table S10), and demonstrated similar complementarity with LSC17 in further stratifying GO response (Fig. S8D; Table S10). Notably, the PC2-34 score remained well correlated with the Primitive vs Mature axis in pediatric AML (r = 0.88) and captured the same survival benefits from GO treatment among these pediatric patients (Fig. S8E-F, Table S10). Together, our data provide a proof of concept that gene expression scores can be readily generated to capture axes of variation in leukemic hierarchy composition and these represent powerful biomarkers of response to non-chemotherapy agents. They are also one of only a few biomarkers that are broadly applicable in both adult and pediatric AML.

### A hierarchy composition framework to guide preclinical studies of AML drugs

We next sought to determine how our leukemic hierarchy framework can be deployed in the context of drug development. Drug candidates are often identified based on reduction in viability of bulk leukemia cells or cell lines, yet this measure lacks critical information pertaining to the subpopulations of cells that are targeted or that persist after treatment. To understand how drug treatment affects cellular composition, we deconvoluted RNA-seq data from 43 datasets in GEO and ArrayExpress with human AML cells sequenced before and after drug treatment (Fig. 5A-B). The changes in cellular composition following drug treatment were visualized in low-dimensional UMAP space and treatments were clustered based on the changes they induced (Fig. 5C-D). Across 153 treatment conditions, 125 resulted in significant changes in cell type composition. Seventy-seven treatment conditions led to a significant increase in PC2, potentially reflecting differentiation, yet most of these treatments resulted in depletion of GMP-like blasts (69%), with fewer treatments depleting the more primitive Quiescent LSPC (30%) or Primed LSPC populations (14%)(Fig. 5D, S10A-B). For example, ATRA induced differentiation predominantly from GMP-like blasts (Fig. 5E). In contrast, differentiation induced by the DHODH inhibitor Brequinar was accompanied by a reduction in Quiescent LSPC abundance, suggesting that this drug may better deplete the stem cell compartment (Fig. 5E).

**Figure 5.**
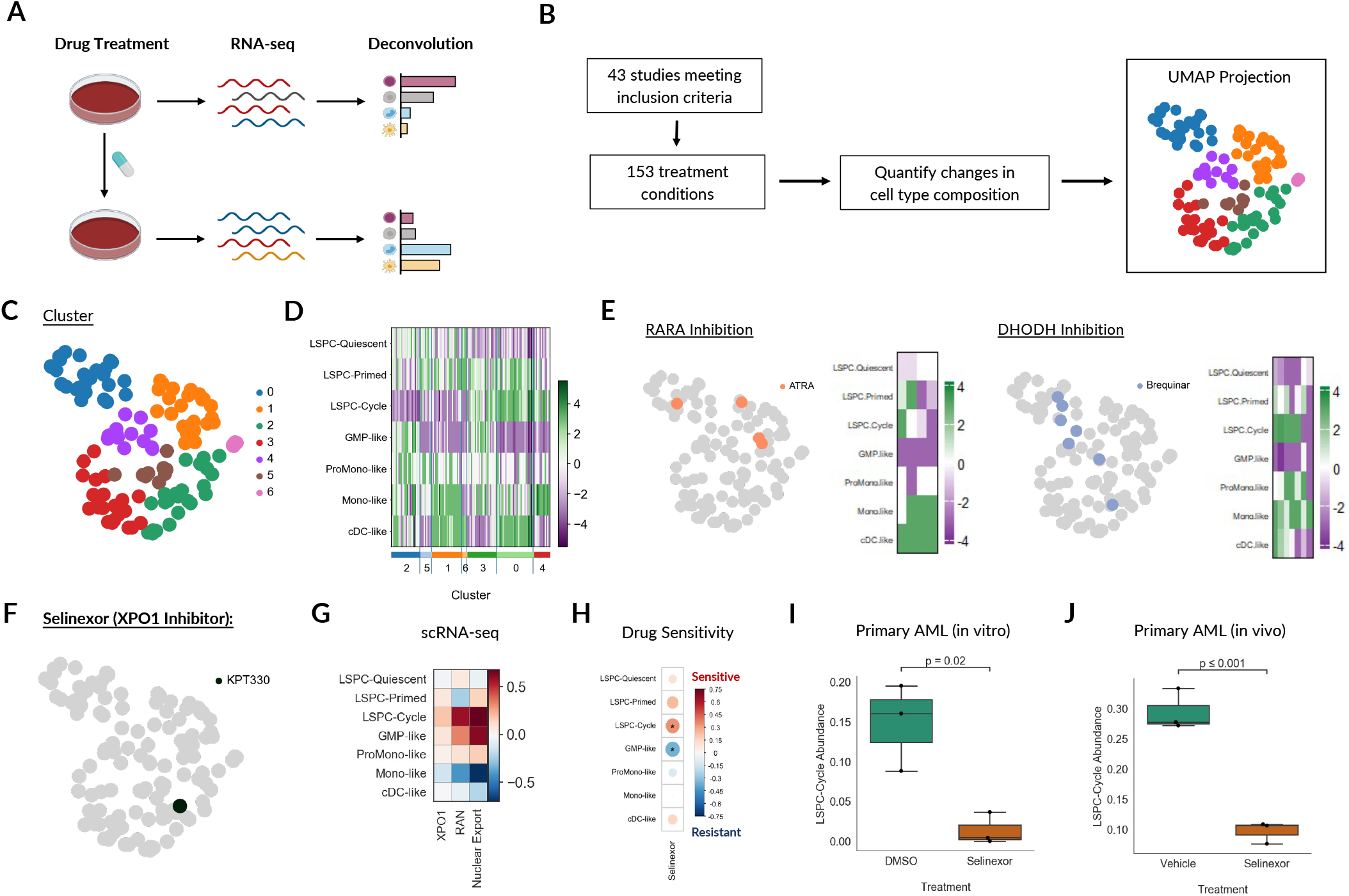
Changes in cellular composition following drug treatment. A) Experimental design of re-analyzed preclinical studies from the literature. Only studies of human AML with RNA-seq available before and after drug treatment were included in order to quantify changes in cell type composition. B) Schematic of re-analysis approach. Changes in the abundance of each cell type were quantified in each treatment condition, and treatments with significant changes in at least one cell type were used as input for dimensionality reduction with UMAP and subsequent clustering. C) Clustering of drug treatments on the basis of changes in cell type composition. D) Heatmap depicting cell type composition changes of drug treatments within each cluster. Purple denotes decreased abundance following treatment and green denotes increased abundance following treatment. E) Examples of the drug treatments targeting specific processes and the changes induced in the abundance of each cell type following treatment. F) Cellular composition changes following *in vitro* Selinexor treatment in NPM1 mutant AMLs from Brunelli et al, 2018. G) Mean expression of XPO1 (the target of Selinexor) and associated genes and pathways in AML blast populations from scRNA-seq. Nuclear export pathway geneset was obtained from GO Biological Pathways. H) Correlation between cell type abundance and *ex vivo* drug sensitivity in BEAT-AML. Correlations with p < 0.05 are marked with an asterisk. I) LSPC-cycle abundance in primary AML samples treated with DMSO control or Selinexor in vitro (Treatment and RNA-seq from Brunelli et al, 2018). J) LSPC-cycle abundance in primary AML samples treated with DMSO control or Selinexor *in vivo* (Treatment from Etchin et al, 2015; RNA-seq from this study).

In some cases, the cell population depleted by a drug corresponded to the expression of the drug target. For example, we analyzed the cellular response to Selinexor, a drug targeting the nuclear export protein XPO1 (Fig. 5F). XPO1 and nuclear export processes were enriched in the Cycling LSPC population at the single-cell level (Fig. 5G), and depletion of this cell population was correlated with *ex vivo* Selinexor sensitivity in the BEAT-AML screen (Fig. 5H). To support this prediction with independent functional evidence, treatment of primary AML samples with Selinexor resulted in depletion of the Cycling LSPC population both *in vitro* ^58^ and *in vivo* ^59^ (Fig. 5I-J) across diverse genetic backgrounds. Together, these data shed light on the changes in cellular composition that follow drug treatment and offer a functionally relevant read-out for prioritizing candidate drugs in preclinical settings.

We next asked how hierarchy information can be utilized in preclinical studies that are more proximal to clinical translation, such as in the context of *in vivo* drug response. To this end we turned to patient-derived xenograft (PDX) response data generated by our group for two drugs: Fedratinib, a JAK2 inhibitor approved for treatment of myeloproliferative neoplasms, and CC-90009 ^60^, an immunomodulatory (IMiD) agent that induces cereblon-mediated degradation of GSPT1 ^61^. A total of 46 independent AML samples were treated in PDX models (n=32 for Fedratinib and n=30 for CC90009) across 658 drug- or vehicle-treated xenografted mice. Deconvoluted RNA-seq profiles from the primary patient samples prior to xenotransplantation were clustered based on hierarchy composition and categorized as Primitive, Intermediate/GMP, or Mature (Fig. 6A-B).

**Figure 6.**
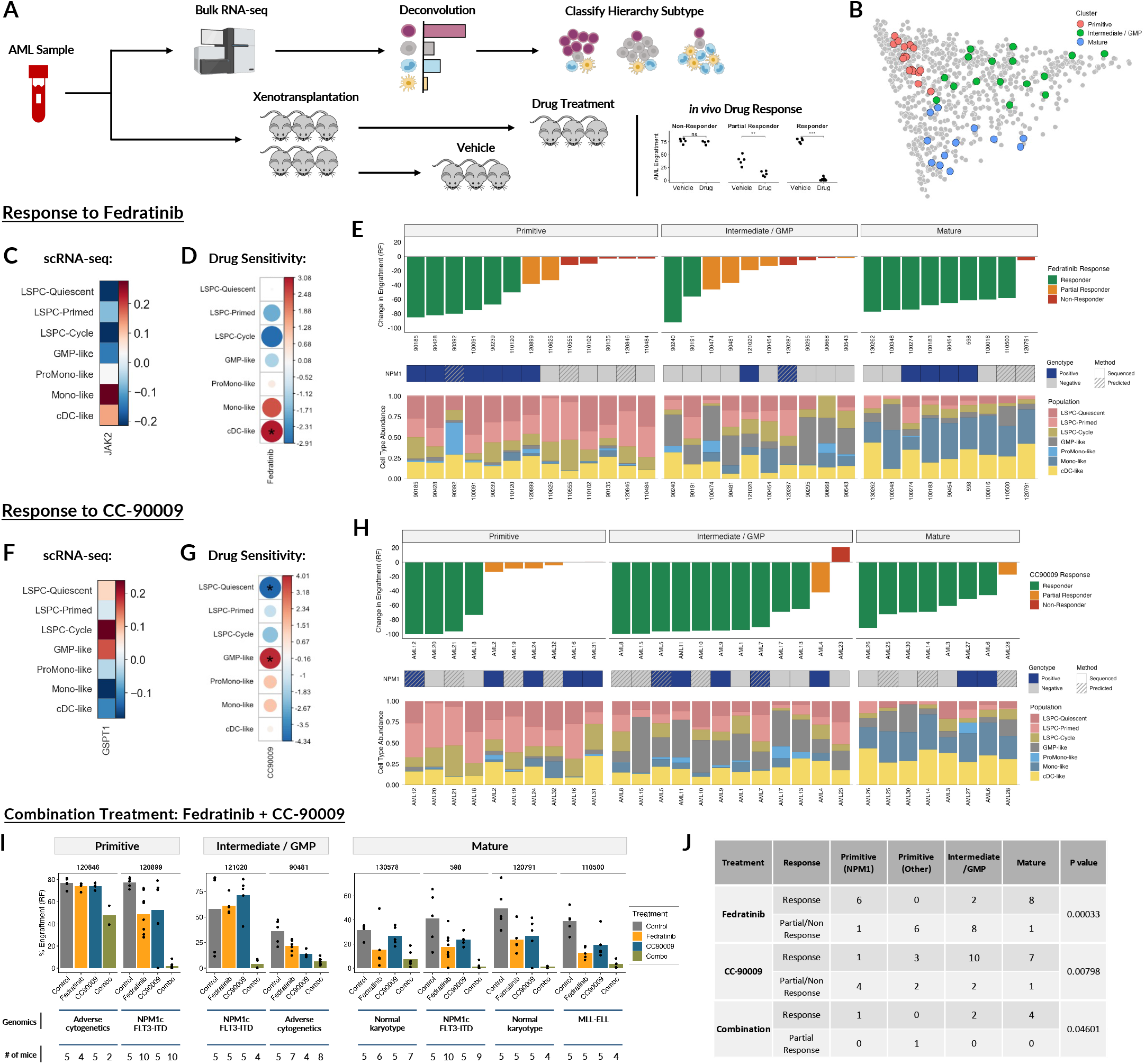
Hierarchy-based stratification predicts *in vivo* response to Fedratinib and CC-90009. A) Experimental design for evaluating the relationship between AML hierarchy composition and drug response in patient-derived xenograft (PDX) models. Response data 658 drug or vehicle-treated treated mice from prior studies (Surka et al, 2021; Chen et al, 2016) were integrated with hierarchy composition data from the primary patient samples, and heterogeneous *in vivo* drug responses were re-analyzed in the context of patient hierarchy subtypes. B) Projected hierarchy composition of primary patient samples prior to *in vivo* drug treatment, categorized by subtype. C) Mean expression of JAK2, the target of Fedratinib, in AML blast populations from scRNA-seq. D) Differences in cell type abundance between full responders and partial / non-responders to Fedratinib. Red depicts enrichment in full responders and blue depicts enrichment in partial / non-responders. Significant differences (p < 0.05) are marked with an asterisk. E) Xenograft responses to Fedratinib, stratified by leukemic hierarchy subtype. Bar plot depicts the mean difference in engraftment in Fedratinib treated mice compared to Vehicle treated mice. Cell type composition of each patient prior to treatment is depicted below each bar. F) Mean expression of GSPT1, the target of CC-90009, in AML blast populations from scRNA-seq. G) Differences in cell type abundance between responders and partial / non-responders to CC-90009, represented as the -log(pvalue). Red depicts enrichment in responders and blue depicts enrichment in partial / non-responders. Significant differences (p < 0.05) are marked with an asterisk. H) Xenograft responses to CC-90009, stratified by leukemic hierarchy subtype. Bar plot depicts the mean difference in engraftment in CC-90009 treated mice compared to Vehicle treated mice. Cell type composition of each patient prior to treatment is depicted below each bar. I) Response to Fedratinib + CC-90009 combination treatment. Patients are stratified by hierarchy and mean engraftment levels are depicted for each treatment condition. J) *in vivo* efficacy of Fedratinib, CC-90009, and Combination treatment of xenografted AML patient samples, stratified by patient hierarchy subtype. Significance was evaluated using a chi-squared test.

The primary target of Fedratinib, JAK2, was predominantly expressed in Mono-like and cDC-like blasts at the single-cell level (Fig. 6C), and these mature blasts were enriched in patient samples that responded well to Fedratinib *in vivo* (Fig. 6D). Subgroup analysis of Fedratinib response showed high efficacy in AMLs with Mature hierarchies (88% response rate), while response rates among other hierarchy subtypes were poor (46% for Primitive and 20% for Intermediate/GMP) (Fig. 6E). CC-90009 targets GSPT1, whose expression is enriched in Cycling LSPC and GMP-like blasts at the single-cell level (Fig. 6F). GMP-like blasts were enriched among responders while Quiescent LSPCs were enriched among partial and non-responders (Fig. 6G). Subgroup analysis showed high CC-90009 efficacy in AMLs with Mature and Intermediate/GMP hierarchies, with 88% and 83% response rates, respectively. In contrast, those with Primitive hierarchies had heterogeneous responses at a rate of 40% (Fig. 6H).

To better understand the heterogeneous responses to Fedratinib and CC-90009 among patient samples, we compared the genomic features of responding and non-responding AML samples for both Fedratinib and CC-90009 treatment conditions. Among Primitive AML hierarchies, *NPM1c* mutations were associated with favorable response to Fedratinib and poor response to CC-90009, while Primitive AMLs lacking *NPM1c* mutation demonstrated favorable response to CC-90009 and poor response to Fedratinib (Fig. 6E,H). Importantly, the association of *NPM1c* signatures with Fedratinib and CC-90009 response among Primitive AMLs did not extend to other hierarchy subtypes. Given the *NPM1c*-based response dichotomy to Fedratinib and CC-90009 among Primitive hierarchies, as well as the sensitivity of Intermediate/GMP hierarchies to CC-90009 and Mature hierarchies to both drugs, we reasoned that a combination of the two drugs may show efficacy against a broader range of samples than either drug alone. To test this hypothesis, we established PDX xenografts from eight AML patients with diverse hierarchy compositions and subjected them to treatment with Fedratinib, CC-90009, both drugs in combination, or vehicle control (median 5 mice/treatment/patient). PDXs from seven of the eight patients tested responded fully to combination treatment with virtual elimination of their leukemic grafts, despite variable responses to single-agent Fedratinib or CC-90009 (Fig. 6I).

Overall, responses to Fedratinib, CC-90009, and combination treatment in PDX models were all significantly associated with hierarchy composition (Fig 6J). These data establish that stratification of AML cases by hierarchy composition facilitates prediction of patients likely to benefit from specific therapies. This also represents a proof of concept for the design of combination regimens through pairing of drugs that show complementarity in their hierarchy-based targeting profiles. Together with the monitoring of post-treatment changes in hierarchy composition, these approaches provide a powerful new paradigm for drug development in AML.

## Discussion

Here we developed a new approach for understanding heterogeneity in AML by characterizing the cellular composition of each patient’s leukemic hierarchy. Analysis of patient-specific variation in hierarchy composition across large cohorts captured and integrated information on genomic profiles, functional stem cell properties, and clinical outcomes within a single classification framework; something that could not be achieved by applying the genomic or stem-cell models alone. Despite the wide diversity of genetic drivers, hierarchy composition could be distilled into four main subtypes, implying convergence in how mutations perturb LSC function and impair hematopoietic differentiation to generate a leukemia cell hierarchy. This new framework provides a means of understanding how different genetic subgroups relate to one another and, more broadly, how genetic alterations relate to LSC properties, enabling a more comprehensive view of biological heterogeneity in AML.

Our analysis of diagnosis/relapse pairs demonstrates the value of longitudinal monitoring of AML hierarchy composition through disease progression. While only a subset of relapsed AML cases are explained by clear patterns of genetic evolution, we show that LSPC expansion constitutes a hallmark of AML relapse following chemotherapy. This has important implications for trial design given that relapsed AML patients, for whom mature and GMP-dominant hierarchies are underrepresented, are often the first patients in which novel therapeutics are evaluated. This mismatch could lead to valuable drugs for patients with such hierarchies being discounted. Our findings also raise an interesting question on the cell types that bear stemness properties. Emerging data from relapse following Venetoclax and Azacitidine treatment shows a loss of phenotypic LSC and the emergence of a promonocytic blast population that may also carry leukemic propagation potential ^62^. Thus an important and unresolved question pertains to the self-renewal capacity of distinct blast populations within these leukemia cell hierarchies. To address this, deeper functional studies of patient samples reflecting specific hierarchy subtypes will be necessary to pinpoint the specific populations that must be targeted to ensure long-term remission.

A large number of targeted therapies are being developed in AML and many investigational therapies are progressing to clinical trials. Our study suggests that biomarkers focused on the composition of each patient’s leukemia cell hierarchy have strong potential to guide the development and selection of these therapies, thereby setting the foundation for a new precision medicine framework in AML. Unexpectedly, prognostic and predictive features were distinct and captured through different axes of variation in hierarchy composition. The Primitive vs GMP axis captured stemness properties including those revealed by LSC17 ^23^ and was highly prognostic. Moreover, other modern prognostic biomarkers ^45–47^ also reflected this axis of variation the same way. Despite capturing stemness and being highly prognostic, drug response to biologically targeted therapies could not be predicted. This indicates that stemness per se is not driving the highly predictive features captured in the Primitive vs Mature axis. Future functional studies are needed to understand the biological mechanism of why this axis drives drug response prediction. Importantly, this axis can be distilled into new gene expression scores including LinClass-7 that have great potential for rapid translation into the clinic. Hierarchy-based classification has clear implications for clinical trial design of investigational drugs: AML samples can now be stratified pre-clinically on the basis of hierarchy composition to identify patient subsets that are most likely to benefit from a single drug or even a predicted combination. Further, for drugs that are already in the clinic such as Venetoclax, Azacitidine, and Gemtuzumab-Ozogamicin where response varies, this stratification could potentially be used to select patients most likely to benefit from these specific treatments. Finally, our study provides a paradigm for translating scRNA-seq information into the clinic that has implications beyond the AML field and could also be applied to other cancers.

## Online Methods

### Patient Samples

All biological samples were collected with informed consent according to procedures approved by the Research Ethics Board of the University Health Network (UHN; REB# 01-0573-C) and viably frozen in the Princess Margaret Hospital (PMH) Leukaemia Bank. No statistical methods were used to predetermine sample size. The investigators were not blinded to allocation during experiments and outcome assessment.

### RNA Sequencing and Pre-Processing

RNA was extracted from bulk peripheral blood mononuclear cells using an RNeasy Micro Kit (Qiagen). Libraries were constructed using SMART-Seq (Clonetec). A paired-end 50-base-pair flow-cell lane Illumina HiSeq 2000 yielded an average of 240 million aligned reads per sample. To align RNA-seq reads from samples used in Selinexor and Fedratinib treatments, Illumina paired-end sequence data were analyzed with BWA/v0.6 alignment software with option (-s) to disable Smith-Waterman alignment. Reads were mapped onto GRCh37-lite reference genome and exon-exon junction reference whose coordinates were defined based on transcript annotations in Ensembl/v59. Reads with mapping quality < 10 were discarded and duplicate reads were tagged using the Picard’s MarkDuplicates program. JAGuaR 2.1 was used to incorporate reads spanning multiple exons into the alignment by introducing large alignment gaps. All transcripts of a given gene were collapsed into a single gene model such that exonic bases were the union of exonic bases that belonged to all known transcripts of the gene. Read counts and subsequently RPKM counts were obtained by counting the fraction of each read that overlapped with an exonic region for that gene. To align RNA-seq reads from functionally annotated LSC fractions, sequence data was aligned against GRCh38 and transcript sequences downloaded from Ensembl build 90 using STAR 2.5.2a. Default parameters were used except for the following: “–chimSegmentMin 12 –chimJunctionOverhangMin 12 – alignSJDBoverhangMin 10 –alignMatesGapMax 100000 –alignIntronMax 100000 – chimSegmentReadGapMax parameter 3 –alignSJstitchMismatchNmax 5 -1 5 5.” Counts were obtained using HTSeq v0.9.1. RNA-seq reads from four AML samples previously profiled by scRNA-seq from van Galen et al ^29^ were aligned to GRCh38 using STAR v2.7.9a and counts were obtained using HTSeq v0.7.2.

### Re-Clustering of Leukemia Stem and Progenitor Cells (LSPCs)

Single-cell RNA-sequencing data from 12 AML patients at diagnosis was obtained from van Galen et al ^29^ (GSE116256). scRNA-seq count data was normalized using the R package ‘scran’ ^63^, log-transformed with an offset value of 1, and scaled. Leukemic cells labeled as “HSC-like” and “Prog-like” (hereafter LSPCs) from the original study were subject to re-analysis using the Self-Assembling Manifolds (SAM) algorithm ^31^. SAM was applied individually to the four patient samples with the highest number of LSPCs (based on a cutoff of >100 HSC-like cells) to assign weights to each gene based on how well they can demarcate emerging transcriptomic states. Feature weights for each gene were averaged across the four samples and subsequently applied to LSPCs from all 12 patients. No batch correction was applied. Using the “scanpy” package ^64^, weighted expression data was subject to dimensionality reduction and neighbourhood detection based on the cell-cell correlation. The diffusion map embedding ^65^ was used for visualization. Leiden clustering ^66^ was performed with a resolution of 0.15 to identify three clusters of LSPCs shared across the patient samples. Re-annotated LSPC labels are included in Table S1.

### Evaluation of LSPC Clustering

To evaluate the new cluster assignments, cell type classifiers were built and evaluated for the new and prior classifications using the R package “SingleCellNet” ^67^. For each classification, scran normalized gene expression values were used as input and 800 cells from each leukemic cell type were used as a training set. For each cell type, paired products of the top 25 genes for each cell type were calculated and the 50 top gene pairs for each cell type were used to train the Random Forest based model with nTrees = 1000. Models trained on the new and prior cell type classification were subsequently evaluated on a held-out dataset of at least 250 remaining cells for each cell type.

### Regulon Analysis and Signature Enrichment

To infer transcription factor (TF) regulon activity in scRNA-seq data, regulon analysis was performed using SCENIC ^68^. The Docker image of pySCENIC was run as per the guidelines from Van de Sande et al ^69^: log-transformed counts from leukemic AML cells were used as the input and candidate transcription factors were identified using a list of human transcription factors from Lambert et al ^70^, with default parameters. To prune putative TF-target links within each regulon using annotations of TF motifs, CisTarget was applied using databases of known human TF motifs annotated at 500bp, 5kb, and 10kb of transcriptional start sites. Drop-out masking was also applied during this step. Enrichment of refined TF regulons was inferred using AUCell, and enrichment scores were scaled for visualization.

### Characterization of scRNA-seq AML Populations

For biological characterization of the re-annotated leukemic cell types, single-cell enrichment scores of hallmark genesets as well as custom genesets from Ng et al (LSC+ AML fractions) ^23^ and Xie et al (S1PR3 overexpression in LT-HSCs) ^33^ were calculated using AUCell ^68^. Cell cycle status was determined using the original annotations from van Galen et al ^29^, in which cell cycle scoring and classification was performed. Shannon diversity of single-cell transcriptomes was calculated from raw count data using the python package “skbio” after down-sampling each cell to 1,000 UMIs.

### Gene Expression Deconvolution

Raw gene expression counts from 13653 cells belonging to any of seven leukemic populations (LSPC-Quiescent, LSPC-Primed, LSPC-Cycle, GMP-like, ProMono-like, Mono-like, cDC-like) or seven non-leukemic immune populations (T, CTL, NK, B, Plasma, and wild-type Monocyte and cDCs) were used as input for signature matrix generation with CIBERSORTx ^71^. Default settings were used with the exception of the minimum expression parameter which was set to 0.25. Deconvolution was performed on TPM-normalized bulk RNA-seq data using S-mode batch correction and Absolute mode. Due to differences in S-mode batch correction performance between the CIBERSORTx web portal and the CIBERSORTx Docker image, we exclusively used the web portal for our analyses. For downstream analysis, the abundance of the seven leukemic populations were normalized to a sum of 1, wherein the score for each population represents the estimated proportion of all leukemic cells. For bulk RNA-seq samples composed entirely of leukemic blasts (cell lines or sorted primary samples), a second signature matrix with seven leukemic populations and no immune populations was used.

For deconvolution with DWLS ^72^, a single-cell signature matrix was generated using MAST ^73^ for each cell type using default settings from the DWLS script. DWLS was then applied to TPM-normalized RNA-seq data using default settings. Deconvolution with Bisque ^74^ was applied to TPM-normalized RNA-seq data following package guidelines and using default settings. Deconvolution with MuSIC ^75^ was applied to TPM-normalized RNA-seq data as per tool guidelines. This was performed in two different ways: direct and recursive. Direct deconvolution involves calculating cell type abundance of each population directly. In order to deal with issues arising from co-linearity, recursive deconvolution was also applied which first calculated the abundance of four groups of cell types: LSPC (LSPC-Quiescent, LSPC-Primed, LSPC-Cycle), GMP (GMP-like), Mature (ProMono, Mono, cDC-like), and Immune (T, B, NK, CTL, Plasma, cDC, Monocyte), and subsequently calculated the abundance of each individual cell type from each group.

### Clinical AML Datasets

Publicly available clinical RNA-seq datasets used for deconvolution analysis are outlined in Table S4. All gene expression data was subject to TPM normalization prior to deconvolution with CIBERSORTx. Clinical and mutational data was extracted from the GDC Data Portal for TCGA (https://portal.gdc.cancer.gov/projects/TCGA-LAML) and from supplemental materials in Tyner *et al* ^40^ for BEAT-AML. For the BEAT-AML cohort, we focused exclusively on pre-treatment samples collected at AML diagnosis (n = 281). For the Leucegene cohort, clinical and mutational annotations were extracted from supplemental materials of 13 papers ^38, 76–87^ and linked based on sample ID.

### Mapping and Clustering AML Hierarchy Composition

To map AML patients based on the composition of their leukemic hierarchies, only deconvolution results pertaining to leukemic AML populations were used. In these cases, estimated abundances from leukemic populations were normalized to 1, such that the value associated with each cell type represents the proportion of total leukemic blasts that it constitutes. Patients from TCGA, BEAT-AML, and Leucegene were used. PCA was performed on the normalized leukemic cell type compositions of these patients. Neighbours were calculated using euclidean distance with a local neighborhood size of 30. To determine the optimal number of clusters, the package “NbClust” was used to calculate 30 clustering metrics for values of k from 2 to 10, and k = 4 was selected by majority rule. Leiden clustering was subsequently performed at a resolution of 0.4 to obtain four hierarchy clusters. Cluster assignments, hierarchy compositions, and genomic annotations for TCGA, BEAT-AML, and Leucegene are included in Table S5.

To project hierarchies onto the reference map from the three AML cohorts (TCGA, BEAT-AML, Leucegene), normalized leukemic cell type abundances from the query dataset was combined with the reference dataset, and batch correction was applied using ComBat ^88^. Following this, the ingest function from scanpy was used to project the batch corrected query dataset onto the principal components of the batch corrected reference dataset and assign cluster labels.

### Hierarchy Classification of Microarray Cohorts

To enumerate patient hierarchy composition from microarray data, we first compared the confidence of CIBERSORTx deconvolution between RNA-seq and microarray data from the TCGA cohort, using the correlation between the original transcriptomes and the synthetic transcriptomes reconstructed from pooling each cell type signature at their estimated frequencies. Deconvolution from RNA-seq achieved the highest correlation (median = 0.95) while deconvolution from Robust Multichip Average (RMA) ^89^ normalized microarray data performed poorly (median correlation = 0.58). However, we found that a single-sample microarray normalization approach (SCAN) ^90^ yielded better deconvolution results (median correlation = 0.80). We next projected the deconvolution results from the microarray samples in TCGA onto the hierarchy map from the reference cohorts using ComBat and the ingest function from scanpy as previously described and compared the assigned cluster labels with the true labels from the corresponding TCGA samples for which RNA-seq data was available. This yielded weighted multi-class accuracies of 0.67 for RMA-normalized deconvolution results and 0.75 for SCAN-normalized deconvolution results. We thus restricted deconvolution of microarray data exclusively to SCAN-normalized data for new cohorts.

Given the comparatively low accuracy (0.75) of our standard approach for hierarchy projection using reference cohorts in the case of microarray data, we employed additional approaches to classify microarray samples from the query cohort as Primitive, Intermediate, Mature, or GMP. A second approach involved projection onto deconvoluted microarray data from TCGA, for which cluster assignments were available from bulk RNA-seq for the same samples. Deconvolution results from the query cohort were batch corrected with the TCGA reference data using ComBat and cluster labels were projected using the ingest function from scanpy. As a third approach, cluster classifiers were trained from the microarray expression data from the TCGA cohort, using the top 10 marker genes for each cluster based on a Wilcoxon test. Microarray expression data from the query cohort was batch corrected with the TCGA reference data using ComBat, and L1-penalty Logistic Regression (L1-LR), L2-penalty Logistic Regression (L2-LR), Support Vector Machine (SVM), K-Nearest Neighbor (KNN), and Random Forest (RF) classifiers were subsequently trained from these marker genes with hyperparameter tuning performed through a grid search with 10-fold cross-validation for each model. For cluster assignment based on gene expression data in the query cohort, the majority vote of all five models was used. To obtain the final cluster assignments, the predictions from all three approaches (projection using deconvoluted RNA-seq reference cohorts, projection using deconvoluted microarray data from TCGA, and gene expression-based classification using microarray data from TCGA) were combined. Within the GSE6891 cohort ^42, 43^, most samples (372/537, 69%) were assigned to a cluster unanimously by all three approaches, while the remaining samples (165/537, 31%) had conflicting assignments between approaches. These ambiguous samples were primarily positioned at the boundary between the Intermediate cluster and other clusters (Primitive, Mature, GMP). These ambiguous cases were reclassified through a KNN approach trained on the 372 high confidence samples using the ingest function from scanpy to obtain the final classifications.

### Classification Benchmarking

To benchmark classification performance for biological phenotypes (e.g. LSC activity, Relapse, Adverse Cytogenetics) as outlined in Fig. S3A, a repeated nested cross-validation approach was employed to obtain high-confidence estimates of model performance. Samples were subject to a 5-fold split (outer cross-validation), wherein each 20% split was used as a held-out set with the remaining 80% used as a training set. Within each split, Logistic Regression or Random Forest classifiers were trained with hyperparameter optimization performed through a grid search with 5-fold internal cross-validation, leading to a total of five separate AUC values. The mean of these five values was calculated as a summary AUC metric, and this nested cross-validation process was repeated for a total of 1000 iterations, with samples being randomly shuffled between each iteration. Together this produces a distribution of 1000 summary AUC metrics, enabling statistical comparisons of model performance across different sets of features. Comparisons were performed through Wilcoxon signed-rank tests, with AUC metrics paired on each iteration, for which the cross-validations splits were the same).

### Clinical and Morphological Correlates

For associations of leukemic and immune cell type abundance with clinical features in TCGA, Pearson correlations were calculated between the absolute abundance of each leukemic and immune population with each clinical feature. Only correlations with uncorrected p < 0.05 were retained. To characterize the hierarchy compositions of distinct FAB morphological classes, we visualized 378 AMLs from TCGA, BEAT-AML, and Leucegene for whom FAB annotations were available. Samples labeled as M5 (without specifying M5A or M5B) were excluded; samples labeled as M6 (n = 4) or M7 (n = 4) were also excluded due to sample size.

### Survival Analysis

Overall survival (OS) was defined as the time from diagnosis until death or last follow-up. Differences in OS between hierarchy classes in each cohort were evaluated using Mantel-Cox Log-Rank tests using the R package “survival”, and survival curves for each cluster were visualized using Kaplan-Meier plots using the R package “survminer”. Univariate and pairwise hazard ratios for each cluster were derived from Cox proportional hazards regression. For combined hazard ratios, individual patient data were pooled and stratified Cox regression was performed with the patient cohort (TCGA, BEAT-AML, GSE6891) set as the stratifier. For multivariate survival meta-analysis, we included covariates that were available across all three cohorts (Cytogenetic Risk, Age, WBC, NPM1 status, and FLT3-ITD status) and performed multivariate stratified cox regression, with patient cohort as the stratifier. We determined whether hierarchy information (e.g. Cluster, PC1, or PC2) adds value in addition to baseline covariates through a likelihood ratio test to assess model improvement after incorporating hierarchy information.

To benchmark the prognostic value of the new LSPC annotation compared to the prior HSC-like/Prog-like annotation, repeated nested cross-validation was performed as described in the classification benchmarking section. Instead of Logistic Regression or Random Forest models, stratified cox regression was performed to predict overall survival from the TCGA and BEAT-AML cohorts, using the abundances of primitive AML populations. L1 (LASSO) or L2 (Ridge) penalties were applied using partial likelihood deviance as the loss function and 5-fold internal cross-validation was performed to identify the optimal lambda value. Rather than AUC, model performance was estimated through the mean likelihood ratio test statistic across the five outer cross-validation splits. This was repeated for 1000 iterations.

To identify the leukemic cell types associated with survival, we performed stratified cox regression on the TCGA and BEAT-AML cohorts using an L1 (LASSO) penalty with partial likelihood deviance loss. Because this process was not repeated over multiple iterations, this score was trained on the full dataset and leave-one-out cross-validation was employed to determine the optimal lambda value. Coefficients for each leukemic population were subsequently used to determine feature importance.

For GSEA analysis of genes ranked by their association with overall survival, only genes that were detected in TCGA and BEAT-AML were evaluated. Univariate Wald tests were performed to evaluate the association with log(TPM+1) normalized expression of each gene with overall survival in each of the TCGA and BEAT-AML cohorts. The Wald test statistics from each cohort were averaged for each gene and used as a rank statistic for GSEA analysis using Stem vs GMP signatures from normal hematopoiesis (top 200 up-regulated and top 200 down-regulated genes from limma differential expression analysis of healthy LT-HSC vs GMP sorted fractions from umbilical cord blood ^33^) and malignant hematopoiesis (top 200 correlated and top 200 anti-correlated genes with PC1).

### Calculation of Prognostic AML Scores

We calculated LSC17 and other prognostic AML scores using log(TPM+1) normalized expression values from TCGA, BEAT-AML, and Leucegene, and normalized microarray expression values from Lee et al ^53^. In cases where specific genes were missing from a dataset, we calculated the score with those genes removed. To ensure high concordance of these partial scores, we calculated the correlation between each partial score and the full score in other datasets to ensure high concordance: we observed a median correlation of r = 0.99 between partial and full scores, with the lowest correlation being r = 0.95. Patients were classified into high and low groups for each score based on a median split within each cohort.

### Mutation Analysis

Cytogenetic and driver mutation annotations from TCGA, BEAT-AML, and Leucegene were used to correlate hierarchy composition with genomic profiles. Mutation combinations between driver mutations were identified and all combinations present in at least 5 patients were retained and visualized along hierarchy axes PC1 and PC2 using the R package “ggridges”. Due to missing variant allele frequency (VAF) information in an appreciable subset of mutation calls from genomic annotations, samples were considered mutated as long as the mutation was called. This analysis was repeated exclusively using mutation calls where VAF > 0.25 to confirm that the observed trends remained the same.

### scRNA-seq Classification in Relapsed AMLs

scRNA-seq profiles of blast cells from 8 relapsed AML patient samples were obtained from Abbas *et al* ^51^. To project these cells onto our cell types defined from diagnostic AML samples from van Galen *et al* ^29^, we used a transfer learning approach implemented through the scANVI ^91^ and scArches ^92^ packages. First, semi-supervised dimensionality reduction was performed with scANVI using unnormalized scRNA-seq data from diagnostic AML samples filtered for 3000 variable genes with malignant cell type annotations and patient batch as a covariate. For scANVI, an initial unsupervised neural network was trained over 500 epochs with patience for early stopping set to 10 epochs, followed by a semi-supervised neural network incorporating cell type annotations that was trained over 200 epochs with a patience of 10 epochs. Transfer learning with scArches was subsequently applied to update the scANVI neural network using scRNA-seq data from the relapsed AML samples, and training was performed over 500 epochs with a patience of 10 epochs. The updated model was subsequently applied to both diagnostic and relapsed AML samples to generate a shared latent representation, and this latent representation was used for further dimensionality reduction with UMAP. For visualization purposes, the diagnostic and relapsed AML data were each downsampled to 10,000 cells.

### Benchmarking Relapse Phenotypes

To benchmark the changes in cellular composition from diagnosis to relapse, we obtained 12,441 gene sets from the MSigDB database corresponding to Hallmark genesets (n = 48), Oncogenic signatures (n = 182), Computationally-derived signatures (n = 667), Chemical and Genetic Perturbations (CGP) from prior Literature (n = 2112), GO Biological Pathways (4400), and previously published Immune Signatures (n = 5024). The relative expression of each signature was scored in each individual diagnosis and relapse sample (n = 88) through GSVA to generate a single-sample enrichment score for each signature. GSVA enrichment scores for each of the MSigDB signatures, alongside the inferred abundance of each leukemic cell type, were compared between diagnosis and relapse through paired t-tests based on the significance of their enrichment at relapse (absolute value of the log10(p)). Each signature was ranked and relapse enrichment of each leukemic subpopulation was subsequently compared against relapse enrichment of each of the MSigDB signatures. Non-parametric Wilcoxon signed-rank tests were also performed for each signature to ensure comparable results.

To benchmark classification performance from using hierarchy information to discriminate between diagnosis-relapse samples, we performed repeated nested cross-validation as outlined in Fig S3A and described in the “Classification Benchmarking” section. This was performed first on individual samples without paired information (n = 88), or on paired patient samples, wherein the changes in cell composition were provided and the classifier was required to identify whether that change in composition corresponded to a transition from diagnosis to relapse (n = 44) or from relapse to diagnosis (n = 44).

### Clonal Evolution Analysis

Clonal analysis of paired diagnosis and relapse samples from four independent cohorts was performed using annotated single nucleotide variant calls derived from targeted sequencing ^48^, whole exome sequencing ^50^, or whole genome sequencing ^18, 49^ data. Genetic clones were identified using PhyloWGS ^93^, selecting the phylogenetic tree with the highest log likelihood (LLH) value. In cases of tied LLH values, the simplest tree with the most representative branching patterns among the top candidates was manually selected. Graphical representations of evolution of genetic clones were depicted using the R package “Fishplot” ^94^ while representations of changes in cell type composition were depicted using the R package “ggAlluvial”.

### Association with Drug Sensitivity

*Ex* vivo drug response in BEAT-AML samples was measured through the Area Under the dose-response Curve (AUC) metric, wherein a low AUC corresponds to sensitivity while a high AUC corresponds to resistance. AUC values were scaled and multiplied by -1 to represent sensitivity in each treatment condition. Pearson correlation was used to measure association between cell type abundance and drug sensitivities, following recommendations from a benchmarking study by Smirnov et al ^95^. Associations were depicted using the R package “corrplot”, and drug sensitivity volcano plots were generated using the R package “EnhancedVolcano”.

### Unsupervised Clustering by Drug Sensitivity

Unsupervised clustering of 30 AML patient samples from Lee et al ^53^ was performed on the basis of their *ex vivo* drug sensitivity values to 159 drugs. Area under the dose response curve (AUC) values for each patient were scaled for each drug, and dimensionality reduction with PCA was applied and neighbours were calculated with a local neighborhood size of 5. Leiden clustering cluster with a resolution of 0.3 was used to determine the final clusters. Differential expression analysis between drug response clusters was performed using limma ^96^ from normalized microarray expression values obtained from GSE107465. The moderated t-statistic for each gene was subsequently used as the rank statistic for GSEA analysis using Stem vs Mature Myeloid signatures from normal hematopoiesis (top 200 up-regulated and top 200 down-regulated genes from limma differential expression analysis of healthy LT-HSC vs Granulocyte/Monocyte sorted fractions from umbilical cord blood ^33^) and malignant hematopoiesis (top 200 correlated and top 200 anti-correlated genes with PC2).

### Derivation of PC2-based Gene Expression Scores

For derivation of PC2-based gene expression scores we used log(CPM+1) normalized gene expression values, which we found to improve model performance during training. To derive the LinClass-7 score, logCPM-normalized expression of 16 genes from the LSC17 assay were used as input features for LASSO regression: DNMT3B, GPR56, NGFRAP1, CD34, DPYSL3, SOCS2, MMRN1, KIAA0125, EMP1, NYNRIN, LAPTM4B, CDK6, AKR1C3, ZBTB46, CPXM1, ARHGAP22. The 17^th^ gene, C19orf77, was excluded due to a lack of expression data in the Leucegene cohort. LASSO regression was performed on negative PC2 (high in Primitive and low in Mature) with leave-one-out cross-validation using the LassoCV function from scikit-learn with a path length of 0.1 to determine the optimal lambda value. Patients from TCGA and Leucegene were combined into a training set, and patients from BEAT-AML were used as a validation set to evaluate the strength of the association between LinClass-7 and PC2.

To train the PC2-34 score, we started with the top 50 correlated and top 50 anti-correlated genes with PC2, based on the average Pearson correlation between the TCGA and Leucegene cohorts. LASSO regression was performed on PC2 with leave-one-out cross-validation to determine the lambda value corresponding to the lowest mean square prediction error. To further reduce the number of features in the model, the largest lambda within one standard error of the lowest root mean square prediction error (RMSE) was selected instead of the lambda directly corresponding to the lowest RMSE. This resulted in a 34-gene score (PC2-34) which was then evaluated in the BEAT-AML validation set.

### Literature Screen for Drug-Treated RNA-seq Datasets

To identify RNA-seq datasets collected from AML samples before and after drug treatment, Applying the search terms “Acute Myeloid Leukemia” and “AML” with the “Homo Sapien” and “RNA-sequencing” flags on Gene Expression Omnibus (GEO) and ArrayExpress, we identified 95 datasets posted before June 17, 2021. From these, 53 were inhibitors that met the inclusion criteria of human AML samples with available RNA-sequencing data collected before and after drug treatment. Datasets with only differential expression results or Bigwig files were excluded. Datasets with less than three samples in each treatment group were also excluded, resulting in a total of 47 datasets included in the final analysis. Detailed information on included datasets is available in Table S12. Each dataset was processed and underwent TPM normalization and deconvolution with CIBERSORTx using a signature matrix of seven leukemic cell types (LSPC-Quiescent, LSPC-Primed, LSPC-Cycle, GMP-like, ProMono-like, Mono-like, cDC-like). For quality control among cell line samples, the deconvolution correlation values from each sample across every dataset were compared and the Jenks natural breaks algorithm was employed to identify cutoffs demarcating low, medium, and high correlation bins. Cell line samples classified as “low-correlation” with a correlation value below 0.437 were excluded from further analysis, leaving 43 datasets spanning 153 treatment conditions.

### Quantifying Hierarchy Composition Changes after Drug Treatment

Relative changes in cell type abundance in each treatment condition were evaluated using Wilcoxon rank-sum tests for technical replicates or Wilcoxon signed-rank tests for biological replicates with paired treatment conditions. For dimensionality reduction with UMAP, we focused exclusively on changes in cell type abundance where the p-value was < 0.05 to emphasize the key changes in cell type composition induced by each drug, resulting in 125 treatment conditions spanning 38 studies. Absolute log p-values were used to represent the magnitude of the shift in cell type abundance, and cell type changes where p > 0.05 were assigned a magnitude of zero. We then applied UMAP with the following parameters (n_neighbors = 13, min_dist = 0.05) to generate the final representation, and Leiden clustering was applied with a resolution of 1. We note that UMAP was selected for visualization rather than PCA because, despite the low number of features, PCA did not adequately capture the variability between clusters. Cell type composition changes for treatment conditions were visualized with the R package “ComplexHeatmap” and are included in Table S13.

### Fedratinib and CC90009 – Hierarchy Classification

Using normalized leukemic cell type composition data for 46 patient samples used for *in vivo* Fedratinib or CC-90009 treatment, dimensionality reduction was performed and clustering was assigned using the Leiden algorithm with a resolution of 0.7, yielding three clusters: Primitive, Mature, Intermediate/GMP. Owing to an under-representation of engrafting samples with GMP hierarchies, we did not attempt to divide the Intermediate/GMP cluster into Intermediate and GMP groups. Samples were subsequently projected on the reference map for visualization and confirmation of cluster assignments.

### Fedratinib and CC90009 – Response Classification

Patient samples were classified into response categories by comparing the relative reduction (RR) of AML engraftment in drug-treated mice versus vehicle-treated mice, as per Galkin et al^97^. RR was calculated as: ((mean % engraftment in control mice) - (mean % engraftment in drug treated mice)) / (mean % engraftment in control mice). Patient sample s were classified as Responders if RR in the injected femur (Right Femur, RF) was >50%, classified as Partial Responders if we observed 20 to 50% RR in the RF or >20% in the non-injected femur (Bone Marrow, BM) only, and classified as Non-Responders if there was no statistically significant difference in engraftment levels between control- and drug-treated mice, or if RR was <20% in both RF and BM.

### Fedratinib and CC90009 – Classification of NPM1 Status in Primary Samples

Patient samples from Princess Margaret Hospital (PMH) in Toronto were classified as NPM1-mutant (NPM1-mut) or NPM1-wildtype (NPM1-wt) based on clinical sequencing results. For patient samples where targeted sequencing data was unavailable, we predicted NPM1 status using gene expression-based classifiers. First, log(TPM+1) normalized RNA-sequencing data from PMH samples and the three reference cohorts (TCGA, BEAT-AML, Leucegene) were combined and batch corrected using ComBat ^88^. Two groups of classifiers were trained: the first group comprised of Logistic Regression (LR), Support Vector Machine (SVM), and Random Forest (RF) classifiers trained on NPM1 status from the reference cohorts and the second group comprised of LR, SVM, and RF classifiers trained on NPM1 status from 46 PMH samples for whom NPM1 genotype was available (out of 88 total samples).

To select features for the first group of classifiers, differential expression (DE) analysis was performed using DESeq2 ^98^ with AML cohort and patient hierarchy cluster as covariates, and DE genes were selected using an absolute log2 fold change threshold > 1 and FDR < 0.01. The top 50 NPM1-mut and top 50 NPM1-wt genes were then used to train LR, SVM, and RF classifiers. Hyperparameter tuning was performed through a grid search with 10-fold cross-validation for each model. The final group-1 classifiers were subsequently evaluated on the 46 PMH samples with NPM1 genotype information. To further account for batch effects between reference cohorts and the PMH cohort, optimal classification thresholds were identified based on the ROC curves for the 46 genotyped PMH samples, yielding final classification accuracies of 0.87 (LR), 0.89 (SVC), and 0.93 (RF). These thresholds were subsequently used for prediction of the remaining 42 PMH samples for whom NPM1 status was missing.

The second group of classifiers was trained directly on the 46 PMH samples for whom NPM1 genotype was available. Given that a subset of the PMH samples did not have raw counts available for DESeq2, we identified NPM1 mutation-specific marker genes through a Wilcoxon rank-sum test with the log(TPM+1) normalized expression. Genes with significant differences in expression between NPM1-mut and NPM1-wt PMH samples at FDR < 0.05 were subsequently filtered to keep genes that were also significantly differentially expressed in the reference cohorts at an absolute log2 fold change threshold > 1 and FDR < 0.01, leaving 63 high confidence genes. LR, SVM, and RF classifiers were subsequently trained from these 63 genes and hyperparameter tuning was performed through a grid search with 10-fold cross-validation for each model. Classification performance was evaluated by 10-fold nested cross-validation with 100 repeats, yielding median accuracies of 0.98 (LR), 1.00 (SVC), and 0.98 (RF). These models were subsequently used to predict NPM1 status in the remaining 42 PMH samples for whom NPM1 status was missing.

For the final prediction of NPM1 status, the classifier with the lowest accuracy (group-1 LR) was excluded and the five remaining classifiers voted on the NPM1 genotype, with the majority vote being assigned as the final prediction. Together, this resulted in high-confidence predictions of NPM1 genotypes for patients in whom targeted sequencing data was not available. Imputed NPM1 genotypes for each of these patients are presented in Figure 6 alongside NPM1 genotypes obtained from sequencing.

### Fedratinib and CC90009 Combination Treatment

NOD.SCID mice were bred and housed in the University Health Network (UHN) animal care facility and all animal experiments were performed in accordance with guidelines approved by the UHN animal care committee. Ten-week-old NOD.SCID mice were irradiated (225cGy) and pretreated with anti-CD122 antibody (200ug per mouse) 24 hours prior to transplantation. Viably frozen mononucleated cells from AML patients were thawed, counted, and intrafemorally injected at the dose of 5 million cells per mouse. At day 21 post-transplantation, treatment of either CC-90009 or Fedratinib alone with vehicle, or in combination, was initiated twice a day for 2 weeks. CC-90009 was given by intraperitoneal (IP) injections at the dose of 2.5mg/kg and Fedratinib was dissolved in 0.5% methylcellulose and orally gavaged at 60mg/kg. Following treatment, levels of AML engraftment were assessed to determine the efficacy of drug treatment against the disease in the mice. Cells collected from the injected right femur, non-injected bone marrow of each individual mouse were stained with human-specific antibodies and evaluated by flow cytometry. Antibodies used for assessment of human AML engraftment include: CD45-APC, CD33-PE-Cy5, CD19-V450, CD34-APC-Cy7, CD15-FITC (BD), CD33-PE-Cy5, and CD14-PE (Beckman Coulter).

## Acknowledgments

We thank C. Jones, A. Tikhonova, N. Iscove, S. Chan, P. van Galen, S. Abelson, B. Haibe-Kains, M. Anders, and all members of the Dick laboratory for valuable feedback on the manuscript. We thank P. Valk and the HOVON-SAKK trial group for providing the clinical annotations associated with GSE6891. A.G.X Zeng is supported by a CIHR Vanier scholarship. J.E. Dick is supported by funds from the: Princess Margaret Cancer Centre Foundation, Ontario Institute for Cancer Research through funding provided by the Government of Ontario, Canadian Institutes for Health Research (RN380110 - 409786), International Development Research Centre Ottawa Canada, Canadian Cancer Society (grant #703212 (end date 2019), #706662 (end date 2025)), Terry Fox New Frontiers Program Project Grant (Project# 1106), University of Toronto’s Medicine by Design initiative with funding from the Canada First Research Excellence Fund, a Canada Research Chair, Princess Margaret Cancer Centre, The Princess Margaret Cancer Foundation, and Ontario Ministry of Health.

## Author contributions

A.G.X.Z., and J.E.D. conceived the project. A.G.X.Z carried out the analysis. S.B. contributed to analyses of post-treatment changes in cell composition. L.J. performed Fedratinib and CC-90009 combination treatments. A.A. and W.C.C. performed RNA-seq extraction and library preparation for LSC fractions and Fedratinib samples, respectively. M.C.-S.-Y. and V.V. performed alignment and preprocessing for LSC fractions and Fredratinib RNA sequence data. H.A.A., N.D., and A.F. provided scRNA-seq data and leukemic cell annotations for relapsed AMLs. P.vG. provided bulk RNA-seq data for AML samples. A.T. analyzed clinical flow data from the PMH Leukemia cohort. M.C., C.P. and H.D. provided GE and clinical data for the ALFA-0701 trial cohort. M.D.M. provided PMH AML samples. J.A.K. and M.D.M. provided clinical annotations for the PMH AML cohort. A.G.X.Z., J.A.K., J.C.Y.W., and J.E.D. interpreted the data. A.G.X.Z. and J.E.D. wrote the paper. J.A.K. and J.C.Y.W. revised the paper.

## Data and Code Availability

Analysis notebooks and all deconvolution results, as well as instructions for applying AML deconvolution, are available at: https://github.com/andygxzeng/AMLHierarchies. Processed RNA-seq data will be uploaded to the Gene Expression Omnibus and raw sequence data will be uploaded to the European Genome-phenome Archive.

## Supplemental Note 1: Benchmarking of deconvolution approaches

### Comparison of Deconvolution Approaches

To identify the deconvolution approach best suited for our application, we compared four distinct methods designed for deconvolution of bulk gene expression from scRNA-seq data, including CIBERSORTx ^1^, DWLS ^2^, Bisque ^3^, and MuSiC ^4^. Each deconvolution approach leveraged single-cell reference profiles from seven leukemic populations (LSPC Quiescent, LSPC Primed, LSPC Cycle, GMP-like, ProMono-like, Mono-like, cDC-like) and seven immune populations (T, B, NK, CTL, Plasma, cDC, Monocyte). In addition to direct deconvolution with MuSIC (MuSIC - direct), we employed a second approach for MuSIC (MuSIC - recursive) wherein the abundance of four groups of cell types LSPC (LSPC Quiescent, LSPC Primed, LSPC Cycle), GMP (GMP-like), Mature (ProMono, Mono, cDC-like), and Immune (T, B, NK, CTL, Plasma, cDC, Monocyte) were first estimated, and the abundance of each individual cell type was subsequently calculated within each group.

We assessed the performance of these deconvolution approaches with regards to three different parameters: 1) accuracy of deconvolution based on parallel bulk RNA-seq and scRNA-seq data collected from the same patients, 2) agreement between deconvolution and flow cytometry for populations with well-defined surface markers, and 3) preservation of associations between cell types including in the setting of collinearity.

### Benchmarking with paired bulk RNA-seq and scRNA-seq data

Benchmarking analyses for scRNA-seq based deconvolution approaches typically rely on deconvolution of pseudo-bulk data, wherein scRNA-seq count data is pooled on a per-patient basis to simulate bulk RNA-seq. However, these artificial transcriptome profiles do not account for the important technical differences between UMI-based scRNA-seq and read-based bulk RNA-seq. Given this, we sought to perform benchmarking analysis using parallel scRNA-seq and bulk RNA-seq from the same patients. To do so, we performed RNA-seq on four diagnostic AML samples which were originally profiled through scRNA-seq with concurrent genotyping in the van Galen 2019 study ^5^. These four patients for which parallel RNA-seq was performed were each representative of one of the four distinct hierarchy subtypes identified in our study (AML916 = Primitive, AML707B = GMP, AML921A = Intermediate, AML556 = Mature).

We first sought to understand the discrepancy between deconvoluted cell type abundance and true cell type abundance from scRNA-seq. Comparing CIBERSORTx (S-mode Batch Correction or No Batch Correction) and other deconvolution approaches (DWLS, Bisque, MuSIC direct, MuSIC recursive), we found that CIBERSORTx with S-mode batch correction had the lowest mean discrepancy between predicted and observed cell type abundances (CIBERSORTx S-mode: 4.5%; DWLS: 5.8%; MuSIC recursive: 6.7%; CIBERSORTx no batch correction: 6.8%; MuSIC direct: 7.2%; Bisque: 8.4%). Across all methods, CIBERSORTx with S-mode batch correction had the fewest high-discrepancy outliers with 73% of the cell type estimates falling within a 5% discrepancy from scRNA-seq (DWLS: 70%; CIBERSORTx no batch correction: 68%; MuSIC recursive: 64%; Bisque: 61%; MuSIC direct: 55%).

We next generated correlation metrics for each patient wherein the relative abundance of all 14 cell types (7 leukemic and 7 immune) from deconvolution were compared against scRNA-seq. Comparing the correlations between deconvolution and scRNA-seq directly for each patient, we found that CIBERSORTx with S-mode batch correction achieves the best overall performance with high correlations within AML916 (Primitive, r = 0.97), AML707B (GMP, r = 0.94), AML921A (Intermediate, r = 0.89). DWLS also achieved high performance (AML916: r = 0.91; AML707B: r = 0.94; AML921A: r = 0.85), while Bisque and MuSIC performed poorly at this task (Fig. S2A). Importantly, correlations within AML556 (Mature) were low for all deconvolution approaches, with the ProMono-like fraction from AML556 frequently being misattributed to other populations (e.g. cDC-like) by deconvolution algorithms (Fig. S2A). Re-analysis of the scRNA-seq data revealed that AML556 ProMono-like cells carried distinct transcriptomic features compared to ProMono-like cells from other patients (Fig. S2B). We hypothesized that this may be underlying the comparatively poor performance of deconvolution using consensus cell type signatures (including ProMono-like) in this particular sample. Re-running CIBERSORTx with cell type signatures specifically derived from AML556 effectively recovers the deconvolution performance in this sample, achieving a correlation of r = 0.86 (p = 0.003) (Fig. S2C).

In summary, deconvolution from leukemic scRNA-seq populations was highly accurate in AML, particularly for CIBERSORTx (most estimates of cell type abundance were within 5% of the true value), but in select patients this performance may drop if the transcriptional profiles of their leukemic populations diverge too much from the consensus (e.g. ProMono-like in AML556).

### Benchmarking with paired bulk RNA-seq and clinical flow cytometry data

As a final benchmark, we considered the association between cell type abundances reported from deconvolution with the surface markers from flow cytometry. While primitive AML cells are known to be inconsistent with regards to immunophenotype, we reasoned that mature myeloid cell types may be more consistent in their presentation of cell surface markers despite their malignant nature. Of the AML patients from PMH for which RNA-seq was performed on their peripheral blood sample, 7 had clinical flow data available from their peripheral blood (PB) sample and 16 had clinical flow data available from their bone marrow (BM) aspirate. We started by comparing the total abundance of mature myeloid AML populations (ProMono-like, Mono-like, and cDC-like) with positivity levels of the pan-myeloid surface marker CD64. Based on deconvolution from CIBERSORTx with S-mode batch correction, we observed a near-perfect correlation of r = 0.98 between PB deconvolution and PB flow (Fig. S2D), and a high correlation of r = 0.90 between PB deconvolution and BM flow, despite variability in composition across tissue sources. We next compared the estimated abundance of the Mono-like population from deconvolution against levels of the monocyte-specific surface marker CD14, observing a correlation of r = 0.99 between PB deconvolution and PB flow (Fig. S2E) and r = 0.80 between PB deconvolution and BM flow. Other deconvolution methods (DWLS, MuSiC, and Bisque) were also accurate, albeit with comparatively lower correlations and variable slopes (Fig. S2D-E).

### Effect of cell type collinearity on deconvolution results

We also sought to assess how each deconvolution approach preserves relationships between each malignant cell type, particularly in the setting of collinearity (for example, with regards to Quiescent LSPC and Primed LSPC). To evaluate this, we captured the correlation between each cell type based on scRNA-seq data across the 12 patients with a dendrogram (Fig. S2F), and compared this with dendrograms generated from deconvolution of TCGA patients ^6^ to understand how well each deconvolution approach preserved the relationship of these cell types when applied to bulk RNA-seq data (Fig. S2G). From the scRNA-seq data, the LSPC populations (particularly Quiescent LSPC and Primed LSPC) were internally correlated while the mature myeloid populations (ProMono-like, Mono-like, cDC-like) were internally correlated. In the context of bulk RNA-seq, deconvolution with DWLS and Bisque resulted in the LSPC populations being anti-correlated and MuSIC in “Direct mode” reported abundances of multiple cell populations as zero across nearly all patients, despite variable abundances of closely associated populations. We found that CIBERSORTx with S-mode batch correction performed best in preserving these cell-cell associations best in the context of bulk RNA-seq.

Taken together, these new benchmarking analyses led to our selection of CIBERSORTx with batch correction as the preferred gene expression deconvolution approach for bulk AML samples from scRNA-seq reference data.

### Deconvolution with leukemic vs healthy reference populations

An alternative approach to deconvolution using leukemic scRNA-seq populations is to deconvolute bulk AML samples using signatures from healthy hematopoietic populations spanning stem, progenitor, and mature. We also deconvoluted bulk AML samples from three cohorts (TCGA, BEAT-AML, and Leucegene) using scRNA-seq signatures from healthy bone marrow sequenced in van Galen et al ^5^, comprised of the following cell types: HSC, Prog, GMP, ProMono, Mono, cDC, T, B, NK, CTL, Plasma. To compare deconvolution confidence between the leukemic and healthy reference signatures, we used the correlation metric from CIBERSORTx which represents the agreement between the original bulk transcriptome and the ‘synthetic’ transcriptome reconstructed by combining the reference signatures of each cell type at their estimated frequencies. In each cohort, deconvolution with leukemic reference signatures yielded consistently higher confidence compared to deconvolution with healthy reference signatures (Fig. S2H).

### Deconvolution of microarray data

Last, we evaluated the performance of deconvolution applied to bulk transcriptomes profiled through microarray compared to bulk RNA-seq. Within the TCGA cohort, a majority of patients were profiled with both microarray and RNA-seq, and deconvolution was applied to both the microarray and RNA-seq profiles from these patients. Based on the correlation between the original and reconstructed transcriptomes for each patient, deconvolution from RNA-seq achieved high performance (median correlation = 0.95) while deconvolution from Robust Multichip Average (RMA) ^7^ normalized microarray data performed poorly (median correlation = 0.58). However, we found that an alternative single-sample normalization approach (SCAN) ^8^ attenuated the loss in deconvolution performance for microarray profiles (median correlation = 0.80) (Fig. S2I).

The differences in deconvolution performance between microarray normalizations were reflected at the level of individual cell types, particularly primitive LSPCs. Estimated cell type abundances were reasonably correlated between RNA-seq and RMA-normalized microarray for Quiescent LSPC (r = 0.68) and Cycling LSPC (r = 0.77) but correlation of Primed LSPC abundance was poor (r = 0.41), with most samples reported as having 0% Primed LSPC abundance from RMA normalized microarray profiles despite variable abundance levels from RNA-seq profiles. In contrast, SCAN normalized microarray profiles did not display this cell type-specific drop out phenomenon, remaining reasonably correlated with RNA-seq estimates for all three LSPC populations (Quiescent LSPC, r = 0.65; Primed LSPC, r = 0.75; Cycling LSPC, r = 0.82) (Fig. S2J). Thus, in the limited cases where deconvolution of microarray profiles was performed, SCAN normalized data was prioritized over RMA-normalization.

### Supplemental Note 2: Comparison of new and prior LSPC annotations for discerning biological phenotypes

Having established that the new LSPC annotations (Quiescent LSPC, Primed LSPC, and Cycling LSPC; derived from direct clustering of primitive AML cells with feature re-weighting through SAM) were better resolved at the single-cell level and led to higher classification accuracy than prior HSC-like and Prog-like annotations from van Galen et al ^5^ (labeled by a random forest classifier trained on normal CD34+CD38- HSCs and CD34+CD38+ Progenitors from healthy bone marrow), we asked whether deconvolution with the new LSPC annotation was also more informative for discerning biologically and clinically relevant phenotypes than deconvolution with the prior HSC/Prog annotation.

To evaluate this, we first asked whether deconvolution using our new cell type annotations (Quiescent, Primed, and Cycling LSPC) can better discriminate between sorted AML fractions with and without functional LSC activity than deconvolution using the prior HSC/Prog annotations. To evaluate this, we built logistic regression and random forest classifiers to predict LSC activity using either the relative abundance of HSC-like and Prog-like populations (prior annotation) or of LSPC populations (new annotation) and measured their performance based on model AUC. To obtain high confidence estimates of model performance, we employed nested cross-validation and repeated this process for 1000 iterations, shuffling the data between each iteration (Fig. S3A).

The logistic regression (LR) classifier and random forest (RF) classifiers trained on HSC-like and Prog-like abundance achieved median AUCs of 0.74 and 0.81 in predicting LSC activity, respectively. Classifiers trained on LSPC abundance achieved median AUCs of 0.77 (LR) and 0.86 (RF), representing a significant increase in performance (p < 2e-16, Fig. S3B). This improvement may in part be reflected by cell type-specific associations with functional engraftment potential. Through deconvolution of sorted AML fractions with the prior HSC/Prog annotation, both HSC-like and Prog-like abundance were associated with engraftment potential (p = 6e-5 and p = 0.0026, respectively). In contrast, deconvolution with the new LSPC annotation revealed that while Quiescent LSPC and Primed LSPC were both associated with functional LSC activity (p = 8e-6 and p = 0.00062, respectively) while Cycling LSPC were not (p = 0.74). This suggests that long-term engraftment potential is restricted to a subset of primitive AML cells, a conclusion that would not be captured by deconvolution with the original HSC/Prog populations. This is likely due to some Quiescent LSPC and Primed LSPC being labeled as Prog-like in the original annotation, potentially contributing to the enrichment of Prog-like cells in LSC+ fractions.

We next compared the prognostic value of the new LSPC populations against the prior HSC-like and Prog-like populations. To evaluate this, both LASSO and Ridge regression models were trained on overall survival in TCGA ^6^ and BEAT-AML ^9^ using either the relative abundance of the three LSPC populations or the two HSC-like and Prog-like populations, stratified by cohort. In these cases, nested cross-validation was performed with 5 inner folds to identify the lambda values with the lowest partial likelihood deviance, and 5 outer folds to estimate model performance based on the mean likelihood ratio test (LRT) statistic. This nested cross-validation process was repeated for a total of 1000 iterations with shuffling of the data prior to each repeat, and the distribution of LRT statistics was compared between the LSPC annotation and HSC/Prog annotation for both L1 (LASSO) and L2 (Ridge) penalties. Prognostic models trained from LSPC abundance consistently and significantly outperformed models trained from HSC-like and Prog-like abundance in the case of both LASSO and Ridge regression (Fig. S3C). This may also be explained by cell type-specific associations with survival: while both HSC-like and Prog-like populations were associated with survival based on stratified cox regression on the TCGA and BEAT-AML cohorts (HSC-like: HR = 3.02, p = 0.028; Prog-like: HR = 2.22, p = 0.025), deconvolution with LSPC populations reveals that associations with survival are restricted to a subset of primitive AML cells. In this context, Quiescent LSPC and Cycling LSPC were significantly associated with survival (Quiescent LSPC: HR = 3.17, p = 0.028; Cycling LSPC: HR = 8.34, p = 0.00035), while Primed LSPC were not (HR = 1.14, p. = 0.849).

To benchmark across additional clinical phenotypes, we compared classifiers trained on HSC/Prog or LSPC abundance in discriminating between diagnostic and relapsed patient samples ^10–13^ (Fig. S3D) as well as between Adverse cytogenetic status and Intermediate / Favorable cytogenetics among patients in the TCGA (Fig. S3E) or BEAT-AML (Fig. S3F) cohorts. In each of these cases, LR and RF classifiers trained on LSPC abundance performed consistently and significantly better in discriminating between these phenotypes than classifiers trained on HSC/Prog abundance. These results are also summarized in Table S2.

Thus, the new LSPC populations capture biological differences specific to primitive AML cells that are not captured in normal hematopoiesis, and the AML-specific biology reflected in these new LSPC populations improves our ability to resolve important biologically and clinically relevant phenotypes compared to the prior HSC-like and Prog-like annotation.

---

**Figure S1.**
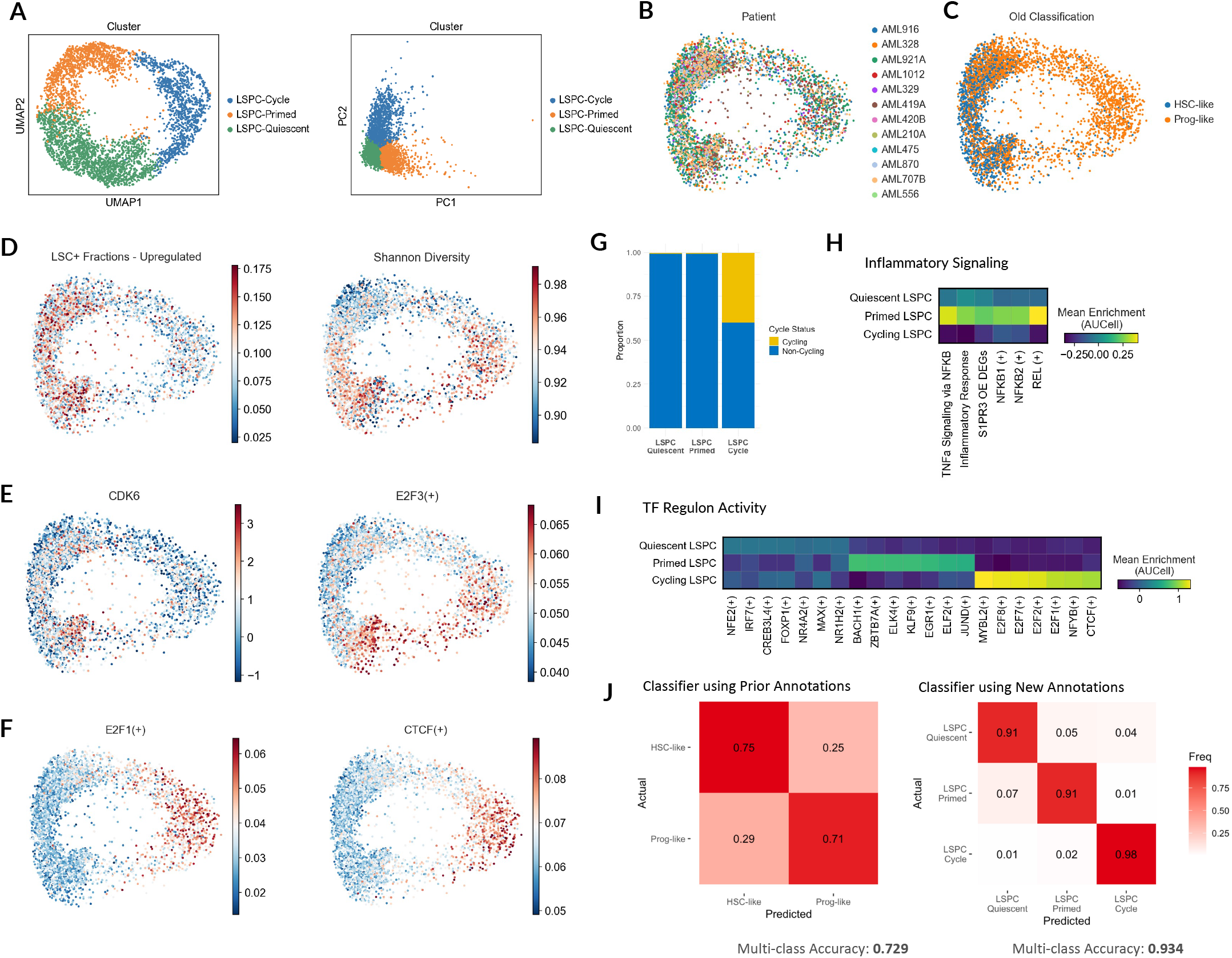
Features of leukemia stem and progenitor cell populations from scRNA-seq. **A)** UMAP and PCA embeddings of AML LSPCs after feature weight derivation with the SAM algorithm. **B-F)** Diffusion map of re-annotated LSPC populations using feature weights from Self-Assembling Manifolds (SAM), depicting: **B)** patient identity, **C)** prior cell type annotation, **D)** enrichment of LSC-specific genes from Ng *et al* (2016) and Shannon Diversity Index, **E)** scaled CDK6 expression and enrichment of the E2F3 regulon, and **F)** enrichment of E2F1 and CTCF regulons. **G)** Cell cycle status of Quiescent, Primed, and Cycling LSPCs. **H)** Enrichment of inflammatory signaling pathways and regulons in LSPCs. **I)** Transcription factor regulon activity, inferred through pySCENIC, specific to each LSPC. **J)** Normalized confusion matrix depicting classifier accuracy of prior and new cell type annotations for primitive AML cells. The classifier was built using SingleCellNet, an Ensemble classifier for scRNA-seq data trained from the top pairs of genes unique to each cell type. 800 cells from each cell type were used for training and 250 were used for validation.

**Figure S2.**
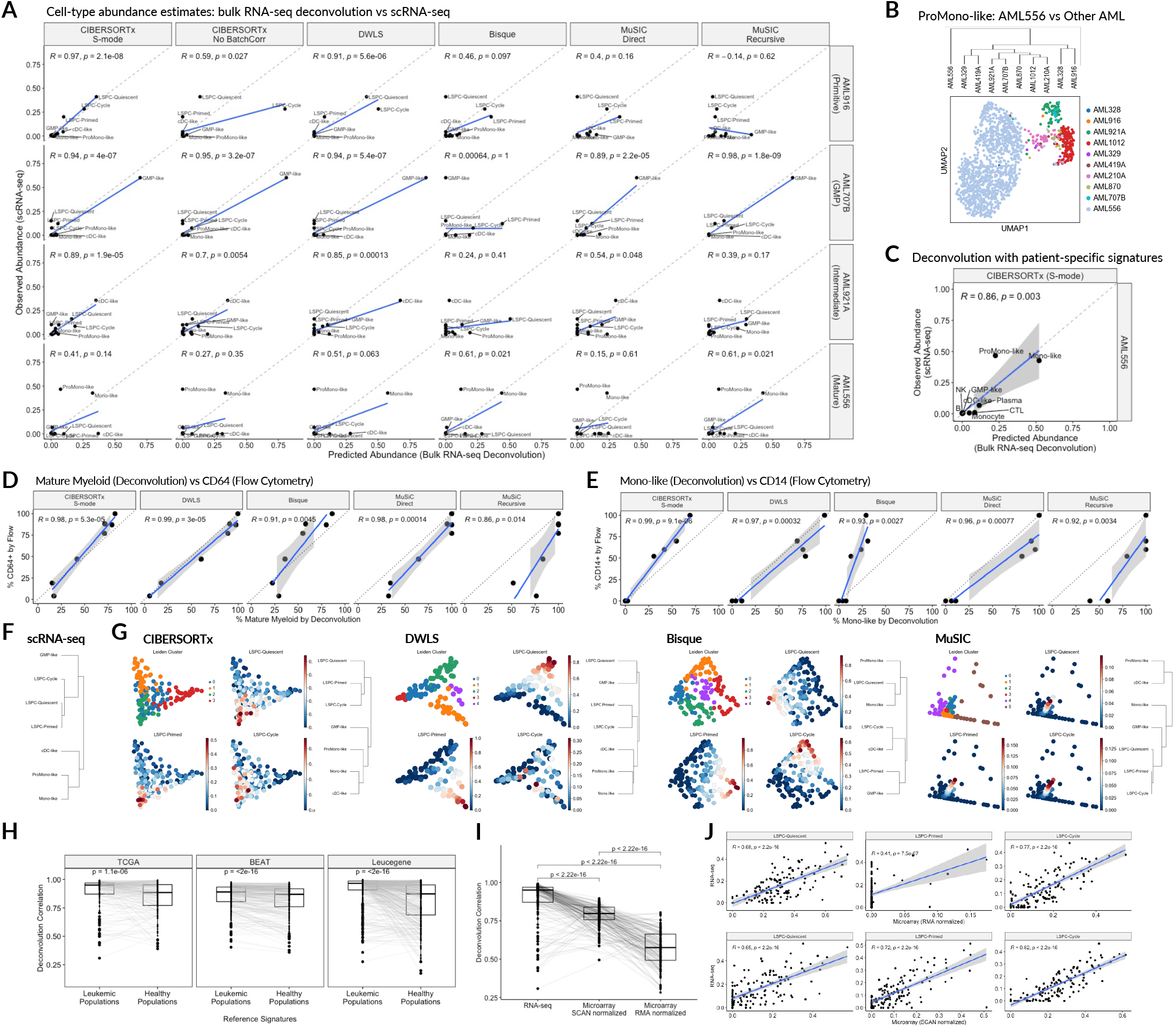
Benchmarking gene expression deconvolution approaches for AML. **A)** Relative abundance of 14 cell types from scRNA-seq compared against inferred abundance from deconvolution of matched bulk RNA-seq data, analyzed by patient. Gene expression deconvolution using CIBERSORTx (S-mode or No Batch Correction), DWLS, Bisque, or MuSIC (direct or recursive) were benchmarked across these samples. **B)** scRNA-seq analyses of ProMono-like cells from AML556 compared to other patients, demonstrating distinct transcriptomic profiles. **C)** Deconvolution performance of CIBERSORTx for AML556 using patient-specific reference signatures. **D-E)** Correlation between deconvolution and clinical flow cytometry for 7 AML patients from the Toronto PMH cohort. Deconvolution using scRNA-seq reference profiles was performed on RNA-seq data and matched with clinical flow cytometry data, both obtained from peripheral blood. **D)** Correlation between total mature myeloid abundance (ProMono-like + Mono-like + cDC-like) from deconvolution with pan-myeloid surface marker CD64. **E)** Correlation between mono-like abundance from deconvolution with monocyte-specific surface marker CD14. **F)** Dendrogram depicting associations between leukemic cell-types across scRNA-seq samples from 12 diagnostic AML patients. **G)** Observed associations between leukemic cell types from deconvolution analysis of 173 patients within the TCGA cohort, depicted for each deconvolution tool. MuSIC Direct was excluded due to multiple cell types having a detection rate of zero in bulk RNA-seq. Correlation between mono-like abundance from deconvolution with monocyte-specific surface marker CD14. **H-I)** Correlation between observed transcriptomic profiles and synthetic transcriptomic profiles reconstructed based on predicted cell-type abundance from CIBERSORTx. Higher correlation suggests greater deconvolution confidence. Comparisons were performed through Wilcoxon signed-rank tests. These correlations are depicted for **H)** Deconvolution of AML cohorts using reference signatures from leukemic populations compared to deconvolution with reference signatures from matched healthy populations, and **I)** RNA-seq compared to microarray from matched TCGA patient samples. Prior to deconvolution, microarray data was normalized through either chip-based (RMA) or single-sample (SCAN) normalization approaches. **J)** Correlation of estimated LSPC abundances between RNA-seq deconvolution and Microarray deconvolution, with either RMA or SCAN normalization, among matched patient samples from the TCGA cohort.

**Figure S3.**
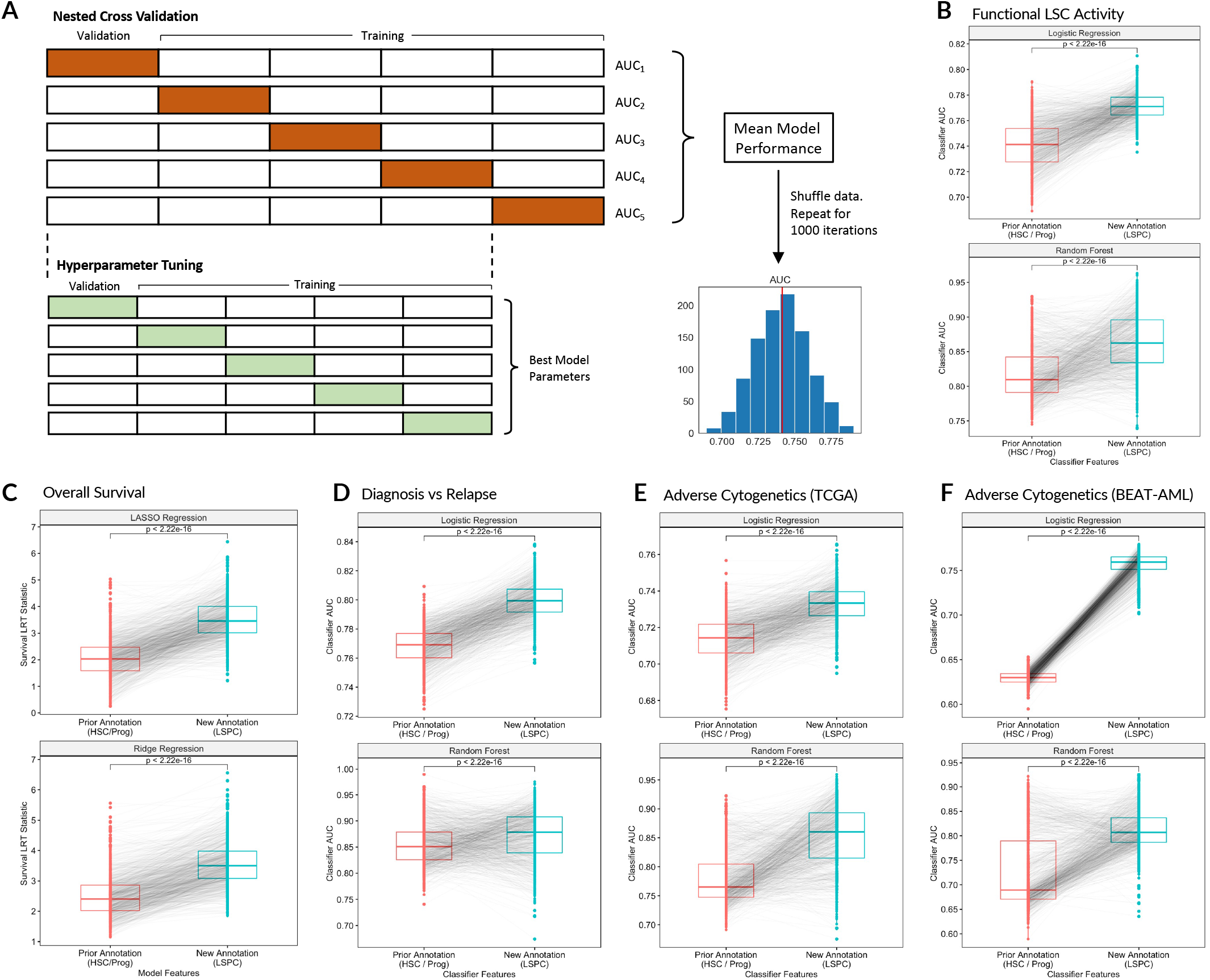
Comparison of new and prior leukemia stem and progenitor cell annotations for discerning biological phenotypes. **A)** Workflow to compare prior (HSC-like and Prog-like) annotations and new (Quiescent LSPC, Primed LSPC, and Cycling LSPC) annotations with regard to their utility in predicting important biological phenotypes in AML. This was measured through the performance of logistic regression and random forest models trained on the relative abundance of these populations. Models were trained using nested cross-validation wherein samples were subject to a 5-fold split, with each split being used to train a unique model with hyperparameter optimization performed by grid search with 5-fold cross validation, and the model AUC for each split was averaged to estimate classifier performance. This nested cross-validation process was repeated over 1000 iterations, with samples being shuffled between each iteration, to generate a distribution of AUC metrics. **B-F)** Model performance for predicting key biological and clinical phenotypes from either HSC-like and Prog-like abundance or Quiescent, Primed, and Cycling LSPC abundance. Performance metrics are paired by iteration, wherein sample order and cross validation splits were identical for each model. **B)** Prediction of functional LSC activity measured by leukemic engraftment from 72 LSC+ fractions and 38 LSC- fractions. **C)** Prediction of overall survival in the TCGA and BEAT-AML cohorts, evaluated through the likelihood ratio statistic from stratified cox regression. In this case LASSO and Ridge regression models were built rather than Logistic Regression and Random Forest classifiers, and these models were trained on splits of the TCGA and BEAT-AML cohorts, stratified by cohort. The repeated nested cross validation approach remained the same. **D)** Prediction of diagnosis vs relapse status from 44 relapsed and 44 diagnostic samples. **E)** Prediction of Adverse cytogenetic status in TCGA from 37 patients with Adverse cytogenetics and 131 patients with Intermediate or Favorable cytogenetics. **F)** Prediction of Adverse cytogenetic status in BEAT-AML from 53 patients with Adverse cytogenetics and 175 patients with Intermediate or Favorable cytogenetics.

**Figure S4.**
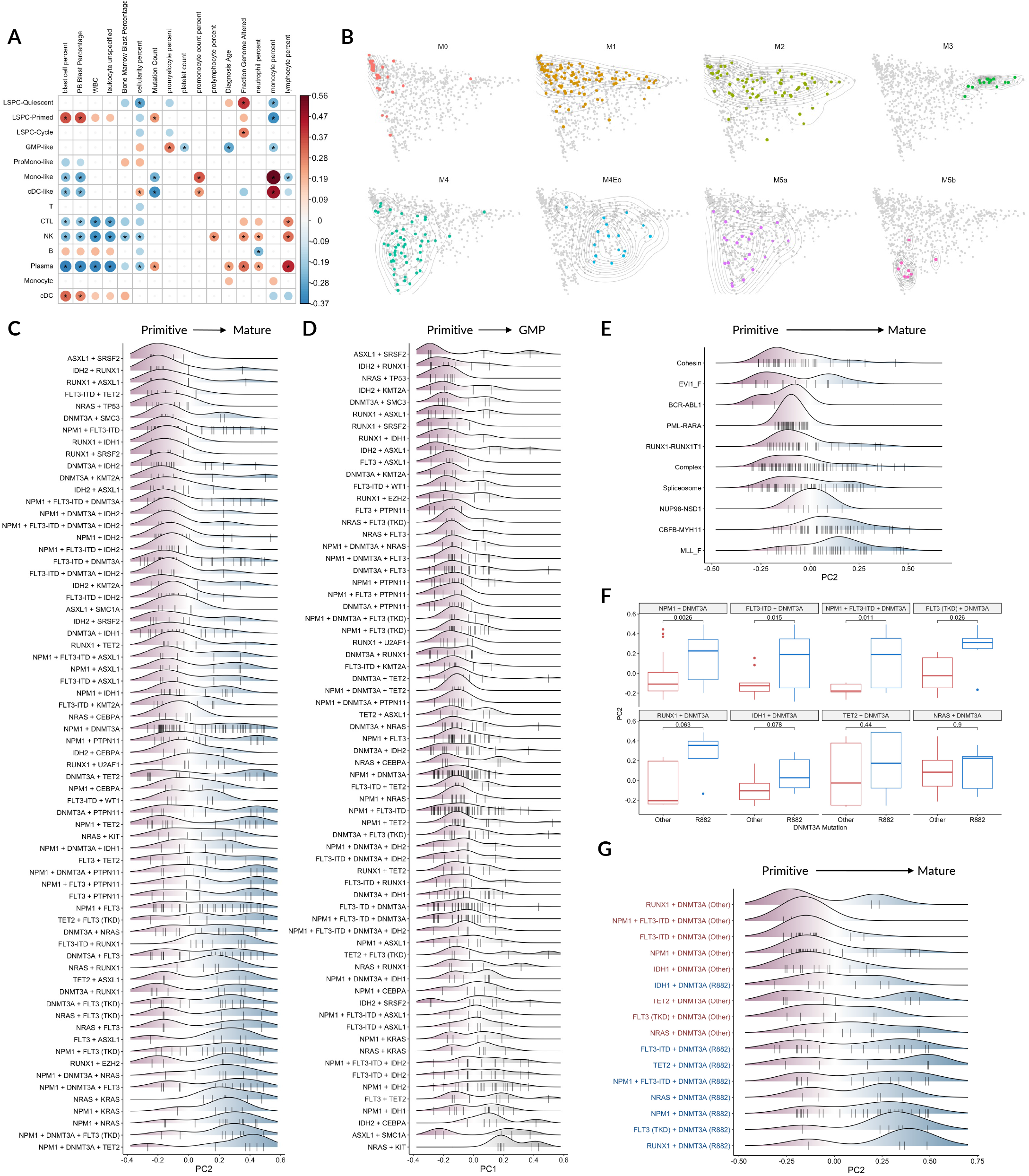
Biological and genomic correlates of AML hierarchies. **A)** Correlation between deconvoluted abundance of leukemic and immune cell types with clinical features in TCGA. Only correlations with p < 0.05 are depicted, and correlations with FDR < 0.05 are noted with an asterisk. **B)** FAB categories from TCGA, BEAT-AML, and Leucegene, projected by cellular hierarchy. **C)** Density plots depicting all mutation combinations along the Primitive vs Mature axis (PC2). **D)** Density plots depicting all mutation combinations along the Primitive vs GMP axis (PC1). **E)** Density plots depicting all cytogenetic alterations along the Primitive vs Mature axis (PC2). **F-G)** Impact of DNMT3A R882 mutations compared to other DNMT3A mutations on leukemic hierarchy organization along the Primitive vs Mature axis (PC2). **F)** Boxplot comparing PC2 of DNMT3A R883 mutant AML compared to other DNMT3A mutations within the context of each mutational combination. **G)** Density plot depicting PC2 of mutational combinations with either DNMT3A R882 or other DNMT3A mutations.

**Figure S5.**
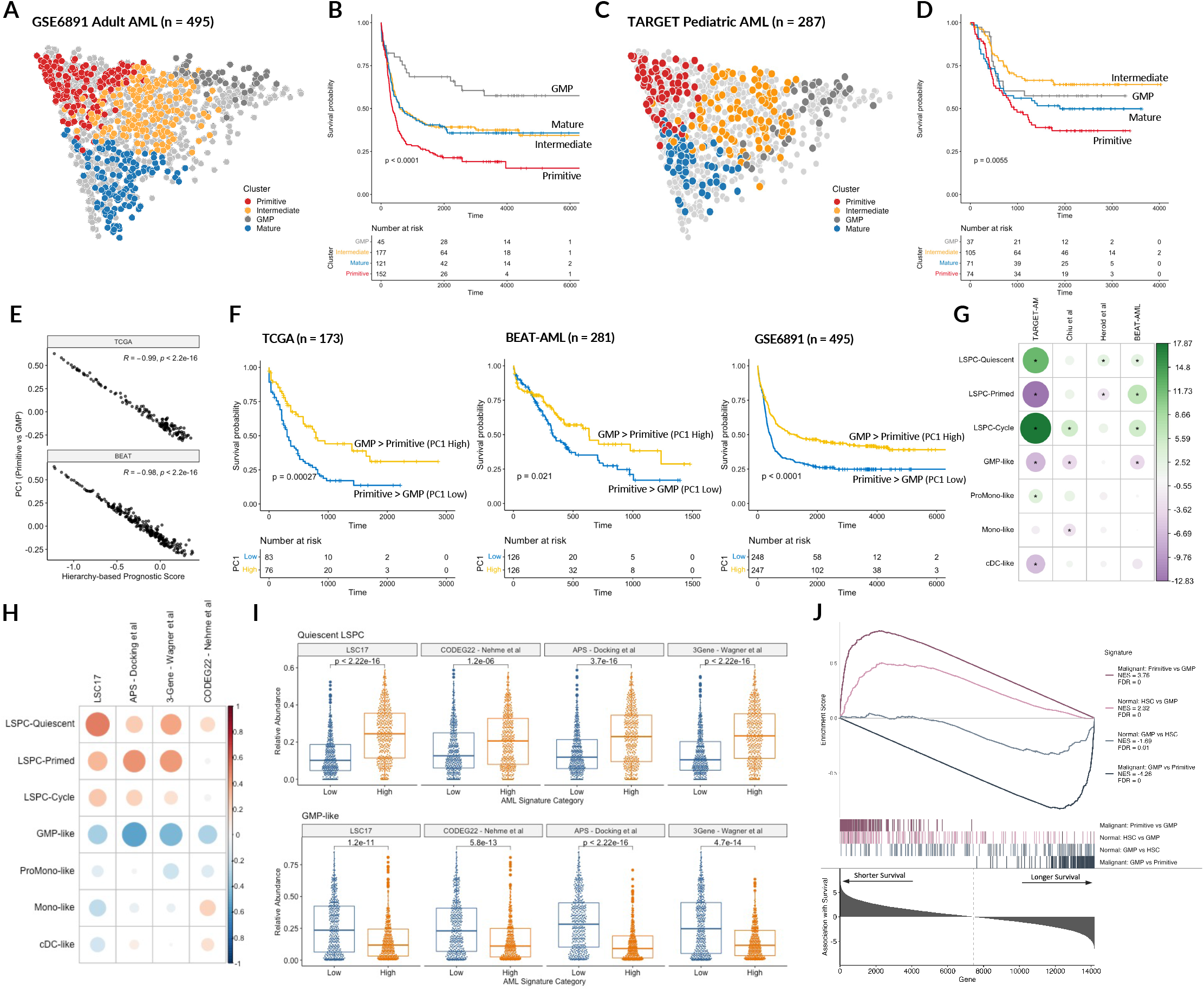
The Primitive vs GMP axis governs AML prognosis. **A)** 495 adult AML patients from GSE6891, projected by hierarchy composition and classified based on the reference cohorts. **B)** Overall survival outcomes across hierarchy subtypes in GSE6891. **C)** 287 pediatric AML patients from the TARGET-AML cohort, projected by hierarchy composition. **D)** Overall survival outcomes across hierarchy subtypes in pediatric AML. **E)** Correlation between a prognostic score trained by regularized cox regression using leukemic cell type abundances with PC1 within the TCGA and BEAT-AML cohorts. **F)** Overall survival outcomes of patients stratified by PC1 within the TCGA, BEAT-AML, and GSE6891 cohorts. **G)** Association between cell-type abundance and induction failure in four independent studies spanning Pediatric AML (Bolouri *et al*, 2018) and Adult AML (Chiu *et al*, 2019; Herold *et al*, 2018; Tyner *et al*, 2018). Green denotes higher relative abundance in induction failure patients compare to patients who achieved complete remission, while purple denotes lower relative abundance in induction failure patients. Differences with a significance of p < 0.10 are noted with an asterisk. **H)** Correlation between four prognostic AML scores with the relative abundance of each leukemic cell type across the TCGA, BEAT-AML, and Leucegene cohorts. **I)** Relative abundance of Quiescent LSPC and GMP-like blasts among AML patients split into high and low risk categories by four prognostic scores. Significance was evaluated through Wilcoxon rank-sum tests. **J)** GSEA analysis with gene signatures derived from the Primitive vs GMP axis in normal and malignant hematopoiesis, performed on genes ranked by univariate associations with overall survival within the TCGA and BEAT-AML cohorts.

**Figure S6.**
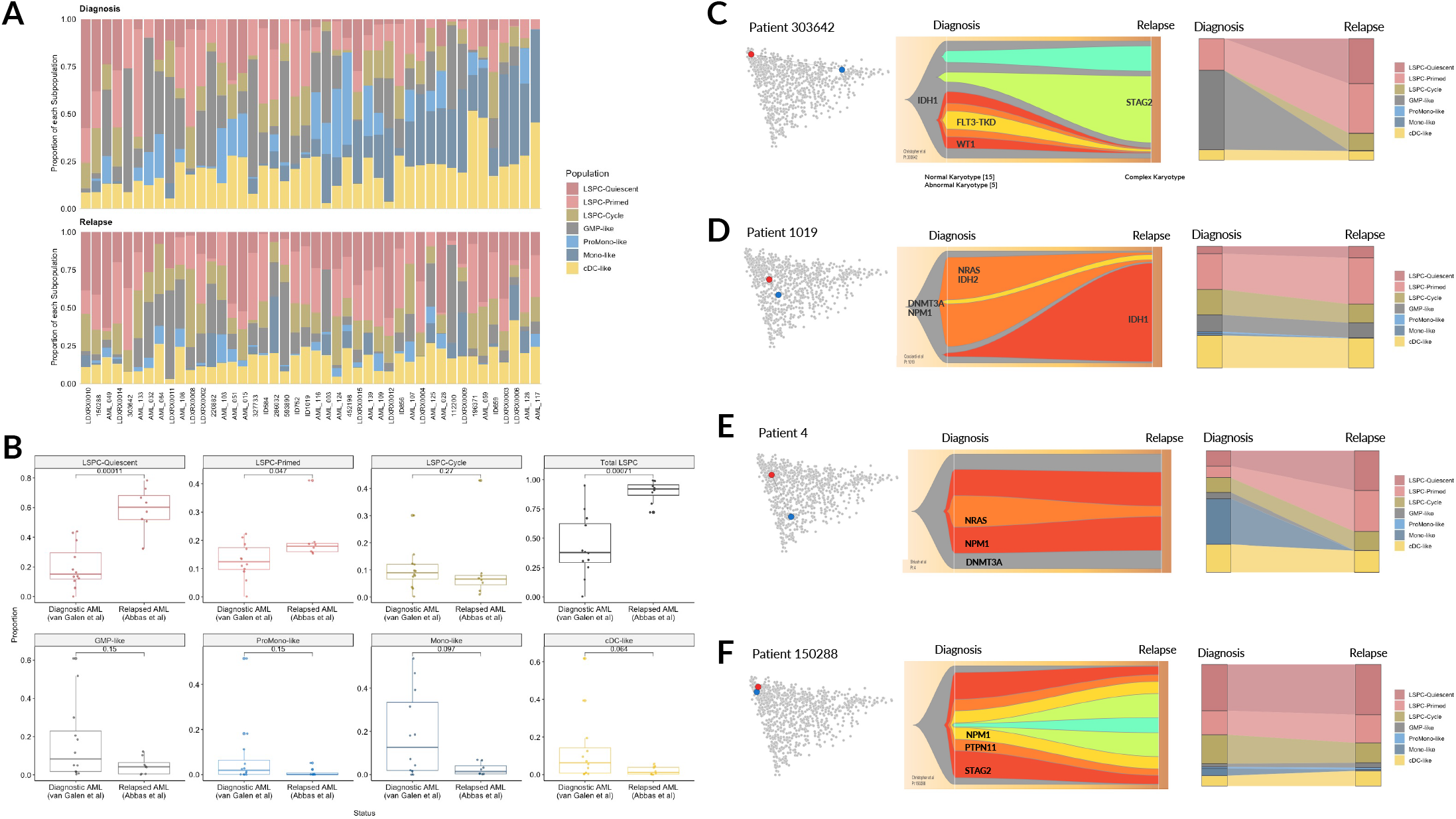
Changes in hierarchy composition at AML relapse. **A)** Hierarchy composition of 44 matched diagnosis and relapse pairs. Top row depicts hierarchy composition at diagnosis while the bottom row depicts hierarchy composition at relapse. Samples from the same patient are aligned vertically. **B)** Relative abundance of each leukemic cell population from scRNA-seq of diagnostic AMLs (van Galen *et al*, 2019) compared with relapsed AMLs (Abbas *et al*, 2021). Statistical significance was evaluated through Wilcoxon rank-sum tests. **C-F)** Evolution of paired diagnosis and relapse AML samples depicted through shifts in cellular hierarchies, evolution of genetic subclones, and changes in cell-type composition. **C)** Patient 303642, in which significant genetic evolution is accompanied by a dramatic shift in cellular hierarchy from GMP to primitive. **D)** Patient 1019, in which replacement of an NRAS and IDH2 positive clone with an IDH1 positive clone is associated with a modest shift in cellular hierarchy. **E)** Patient 4, in which a loss of monocytic blasts is accompanied by a modest decrease in the the size of an NRAS bearing clone. **F)** Patient 150288, in which extensive linear genetic evolution is not associated with any appreciable change in cell type composition.

**Figure S7.**
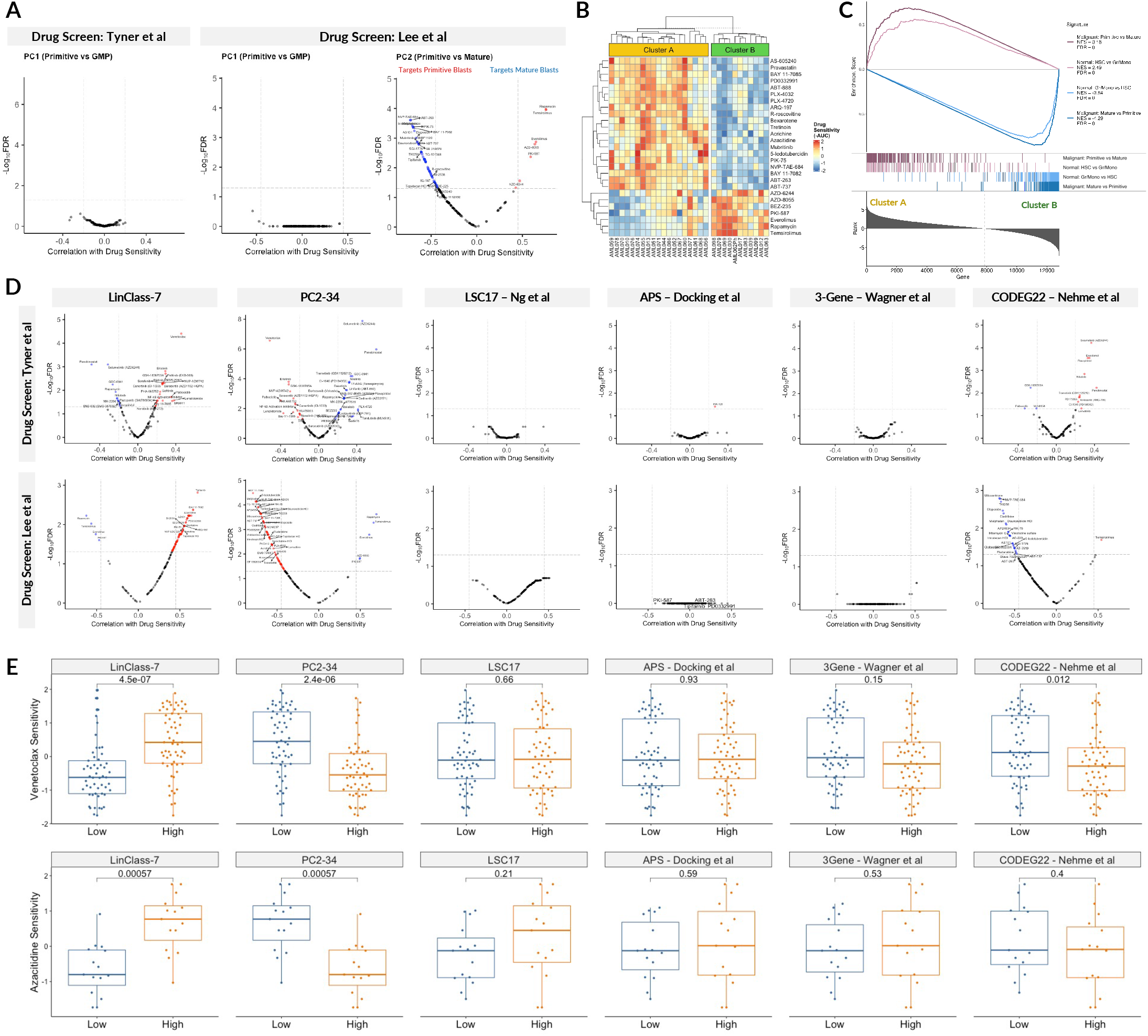
The Primitive vs Mature axis governs ex vivo drug sensitivity. **A)** Volcano plot depicting associations between Primitive vs GMP axis (PC1) and *ex vivo* drug sensitivity from the BEAT-AML (Tyner *et* al 2018) drug screen, and between PC1 and PC2 and *ex vivo* drug sensitivity from Lee et al (2018). **B)** Unsupervised clustering of 30 primary AML patients from Lee *et al* (2018) on the basis of *ex vivo* sensitivity to 159 drugs. Drug sensitivity values (scaled negative AUC) are depicted for the top drugs corresponding to each patient cluster. **C)** GSEA analysis with gene signatures derived from the Primitive vs Mature axis in normal and malignant hematopoiesis, performed on genes ranked by differential expression between the two drug response clusters from (B). **D)** Correlations of AML gene expression scores with *ex vivo* drug sensitivities from two drug screens (Tyner *et al* 2018, Lee *et al* 2018) performed on primary AML samples. **E)** *Ex vivo* drug sensitivity to Venetoclax and Azacytidine of primary patient samples grouped into “High” or “Low” based on median splits of patient scores for each AML gene expression score. Significance was evaluated through Wilcoxon rank-sum tests

**Figure S8.**
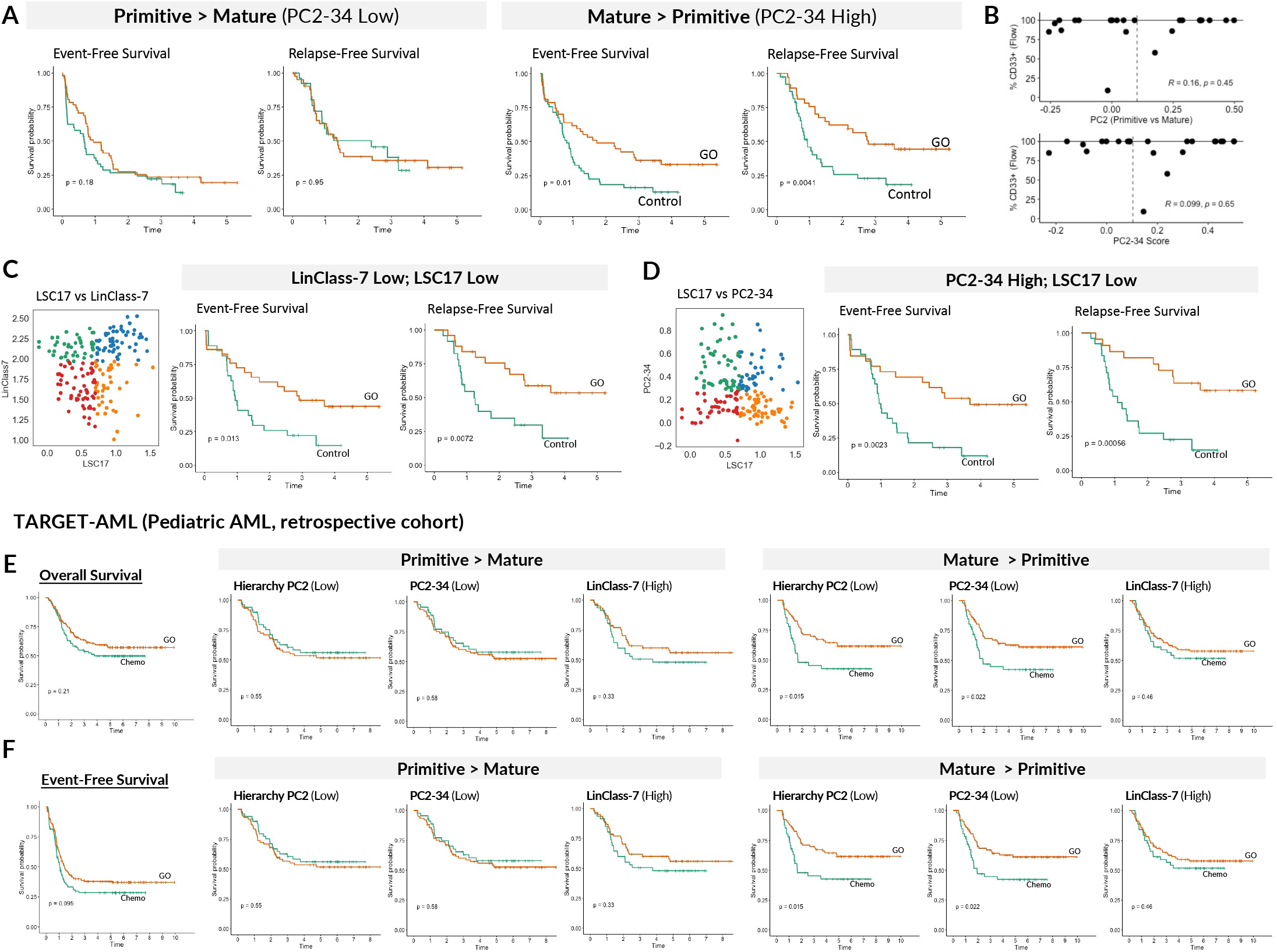
The Primitive vs Mature axis predicts clinical benefit from Gemtuzumab-Ozogamicin in both adult and pediatric AML. **A)** Subgroup analysis of ALFA-0701 after stratification by PC2-34. Event-free and Relapse-free survival curves comparing chemotherapy only (Control arm) against chemotherapy + Gemtuzumab-Ozogamicin (GO arm). **B)** Lack of correlation between CD33 levels by flow cytometry and Primitive vs Mature axis (PC2 and PC2-34 score), evaluated across 23 Toronto PMH AML patients for which both RNA-seq and clinical flow information was available. **C)** Stratification of ALFA-0701 patients on the basis of both LSC17 and LinClass-7. Event-free and Relapse-free survival for the LinClass-7 Low (Mature > Primitive) and LSC17 low subgroup is depicted as this was the only group to derive significant benefit from GO treatment. **D)** Stratification of ALFA-0701 patients on the basis of both LSC17 and PC2-34. Event-free and Relapse-free survival for the PC2-34 High (Mature > Primitive) and LSC17 low subgroup is depicted as this was the only group to derive significant benefit from GO treatment. **E-F)** Subgroup analysis of pediatric AML patients treated with GO or Chemo, stratified by the PC2 Primitive vs Mature axis and related gene expression scores. Outcomes are depicted for both overall survival (E) and event-free survival (F).

**Figure S9.**
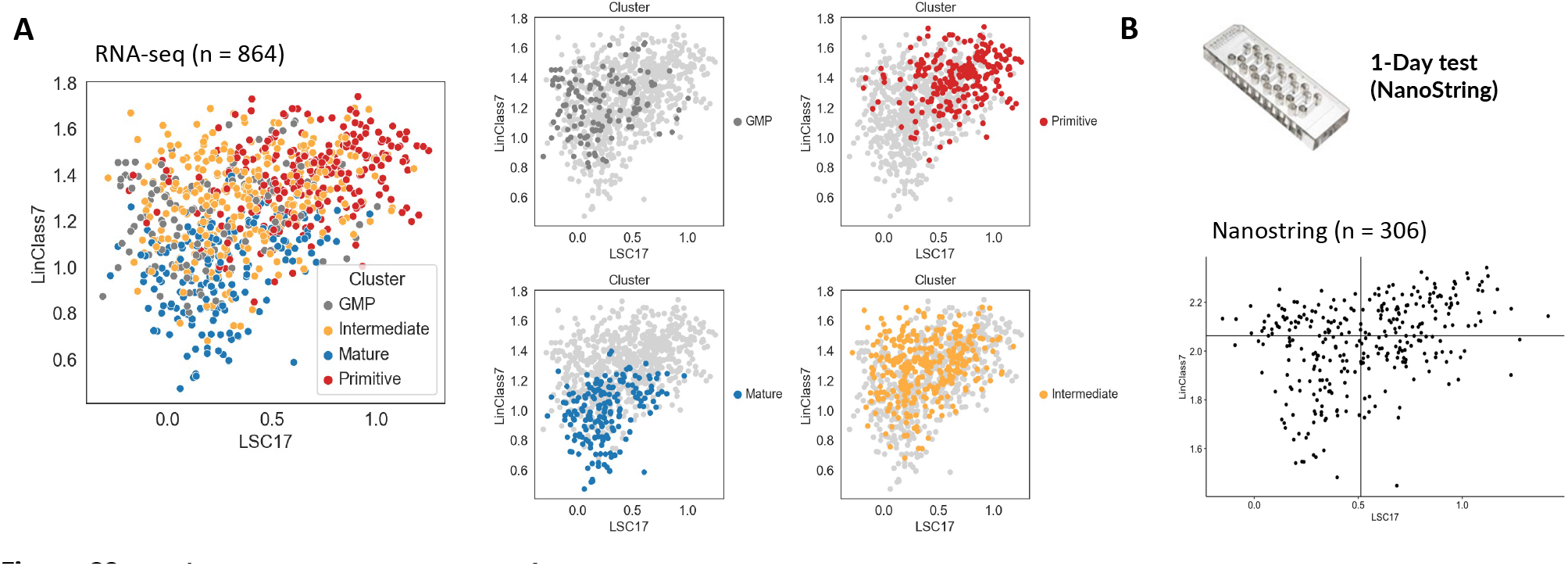
LinClass-7 as a companion score for LSC17. **A**) LSC17 and LinClass-7 scores of 864 AML patients by RNA-seq. Patients belonging to each hierarchy subtypes (Primitive, Intermediate, GMP, Mature) are also depicted. (**B**) LSC17 and LinClass-7 scores measured through a 17-gene NanoString assay. Normalized NanoString-derived LSC17 and LinClass-7 scores from 306 Toronto PMH patients from Ng *et al* (2016) are depicted.

**Figure S10.**
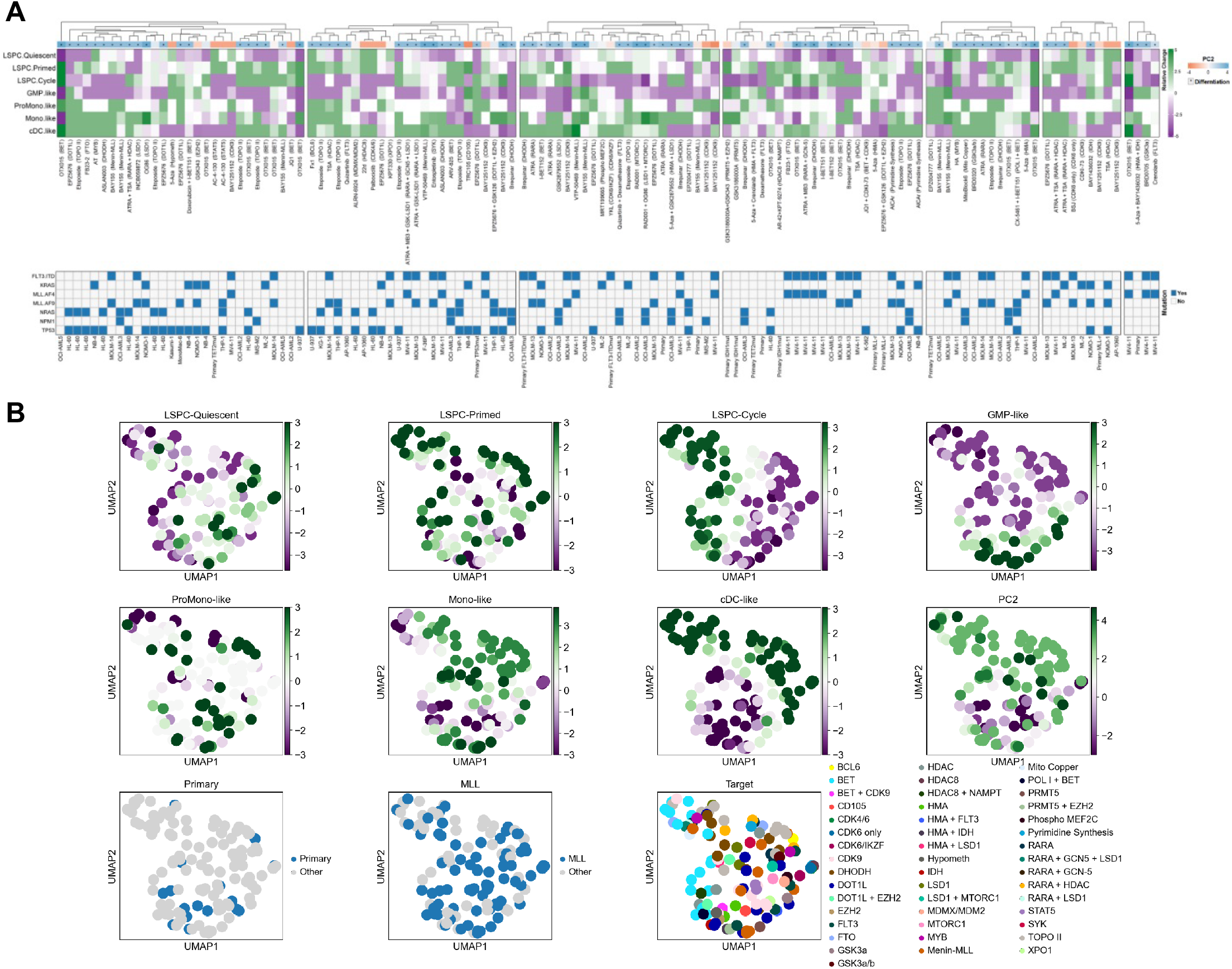
Literature screen to identify treatment-induced changes in cellular composition. **A)** ComplexHeatmap depicting changes in cell type composition following drug treatment from preclinical studies in human AML. Green depicts an increase in cell type abundance and purple depicts an decrease in cell type abundance. Each treatment is labeled with its target(s) in parentheses. Changes in PC2 (Primitive vs Mature) are depicted above the heatmap, and candidate differentiation drugs (increase in PC2 with uncorrected p-value < 0.05) are denoted with an asterisk. AML sample type and key genomic characteristics are also depicted for each treatment condition. **B)** UMAP coordinates for each drug treatment condition depicting changes to each cell type, as well as tissue source (Primary vs Cell Line), MLL translocation status, and drug target.

